# Evolved bacterial formate assimilation is likely potentiated by a rudimentary CO_2_ concentrating mechanism

**DOI:** 10.1101/2025.05.05.652280

**Authors:** Brittany E. Mazny, Breah LaSarre, Alekhya M. Govindaraju, James B. McKinlay

**Affiliations:** Department of Biology, Indiana University, Bloomington, IN

## Abstract

Formate is a single-carbon compound that is challenging to assimilate, including when assimilation involves a CO_2_ intermediate that can diffuse from the cell. Mutations that overcome such challenges can be identified through adaptive laboratory evolution. We evolved the anoxygenic phototrophic bacterium *Rhodopseudomonas palustris* to use formate as the sole carbon source. Through gene deletions, we determined that formate is assimilated via oxidation to CO_2_ by formate dehydrogenase followed by CO_2_ fixation by the Calvin cycle. However, this pathway had no clear link to three genes that were commonly mutated in evolved isolates: (i) *ribB*, a flavin synthesis enzyme, (ii) *ppsR2*, a repressor of light-harvesting genes, and (iii) RPA0893, a regulator of unknown function. A RibB mutation was necessary and sufficient for formate assimilation and improved formate oxidation. PpsR2 mutations occurred early, facilitated formate assimilation, and caused elevated pigmentation. Pigment production generates CO_2_ and alkaline conditions that, along with intracellular chromatophore membranes, could represent a rudimentary but important CO_2_-concentrating mechanism. RPA0983 mutations emerged late and facilitated formate assimilation. RPA0983’s proximity to a CO_2_-liberating pigment synthesis gene suggests a similar effect as PpsR2 mutations. Our findings reveal unintuitive pathway intersections that could have broad implications for formate and CO_2_-utilizing organisms.

## INTRODUCTION

Formate is a single-carbon compound found in environments ranging from the atmosphere (1) to the sea floor (2). Some bacteria can assimilate formate via direct condensation onto a tetrahydrofolate intermediate (Fig S1). This indirect mechanism overcomes an intrinsic difficulty in activating formate for electrophilic or nucleophilic attack (3). Some bacteria assimilate formate indirectly by first oxidizing it to CO_2_, followed by CO_2_-fixation via the Calvin-Benson-Bassham (CBB) cycle (3, 4). A possibly overlooked challenge of the indirect mechanism is the potential loss of the CO_2_ intermediate, due to its high membrane permeability.

How microbes evolved to overcome challenges in formate assimilation can conceivably be inferred from comparative genomics (5, 6). In general, the evolution of novel traits can involve gene amplification (7), horizontal gene transfer (8), gene duplication and divergence of function (9), activation of cryptic genes (10, 11), and mutations that hone promiscuous enzyme activities towards a new function (12). Inferring how genomic changes leads to novel metabolic traits requires prior knowledge of gene function or connectivity. However, novel traits can also emerge by less intuitive means.

Adaptive laboratory evolution leverages the large populations and the short generation times of microbes to monitor genomic changes underlying novel traits in near-real time (5, 13, 14). Adaptive laboratory evolution experiments have led to novel utilization of citrate (7) and propanediol (11) by *Escherichia coli*, glucose by *Shewanella oneidensis* (15), lignin dimers by *Novosphingobium aromaticivorans* (16, 17), and a variety of carbon sources by *Pseudomonas aeruginosa* (6). When genetically tractable bacteria are evolved, researchers can connect genotype to phenotype by engineering mutations into the ancestor and repairing mutations in evolved strains. Such an approach can also inform on the relative contribution to a phenotype. For evolved citrate-utilization by *E. coli*, the trait evolved via (i) potentiating mutations that established a genetic landscape favorable for an (ii) actualizing mutation that was directly responsible for the novel trait, followed by (iii) refining mutations that improved the trait (7).

*Rhodopseudomonas palustris* is a purple phototrophic bacterium that is renowned for its metabolic versatility. However, the type strain CGA0092 (18, 19) cannot use formate as the sole carbon source. CGA0092 is missing the formate-tetrahydrofolate ligase of the direct assimilation pathway. Related bacteria can indirectly assimilate formate via formate dehydrogenase (FDH) or formate hydrogen-lyase followed by CO_2_ fixation via CBB cycle, in some cases requiring exogenous CO_2_ (20-22). *R. palustris* CGA0092 can use CO_2,_ and it has an annotated formate dehydrogenase (FDH; *fdsGBA*, RPA0732-4; Fig 1). Thus, CGA0092 could conceivably evolve formate assimilation, but it is not known to have CO_2_ concentrating mechanisms like cyanobacterial carboxysomes (23) or algal pyrenoids (24). Although CO_2_ concentrating mechanisms are best known to favor CO_2_ fixation over wasteful oxygenation by the CBB enzyme ribulose 1,5-bisphosphate carboxylase (Rubisco), a CO_2_ concentrating mechanism might also be necessary to capture CO_2_ from FDH before it diffuses from the cell.

**Fig 1.**
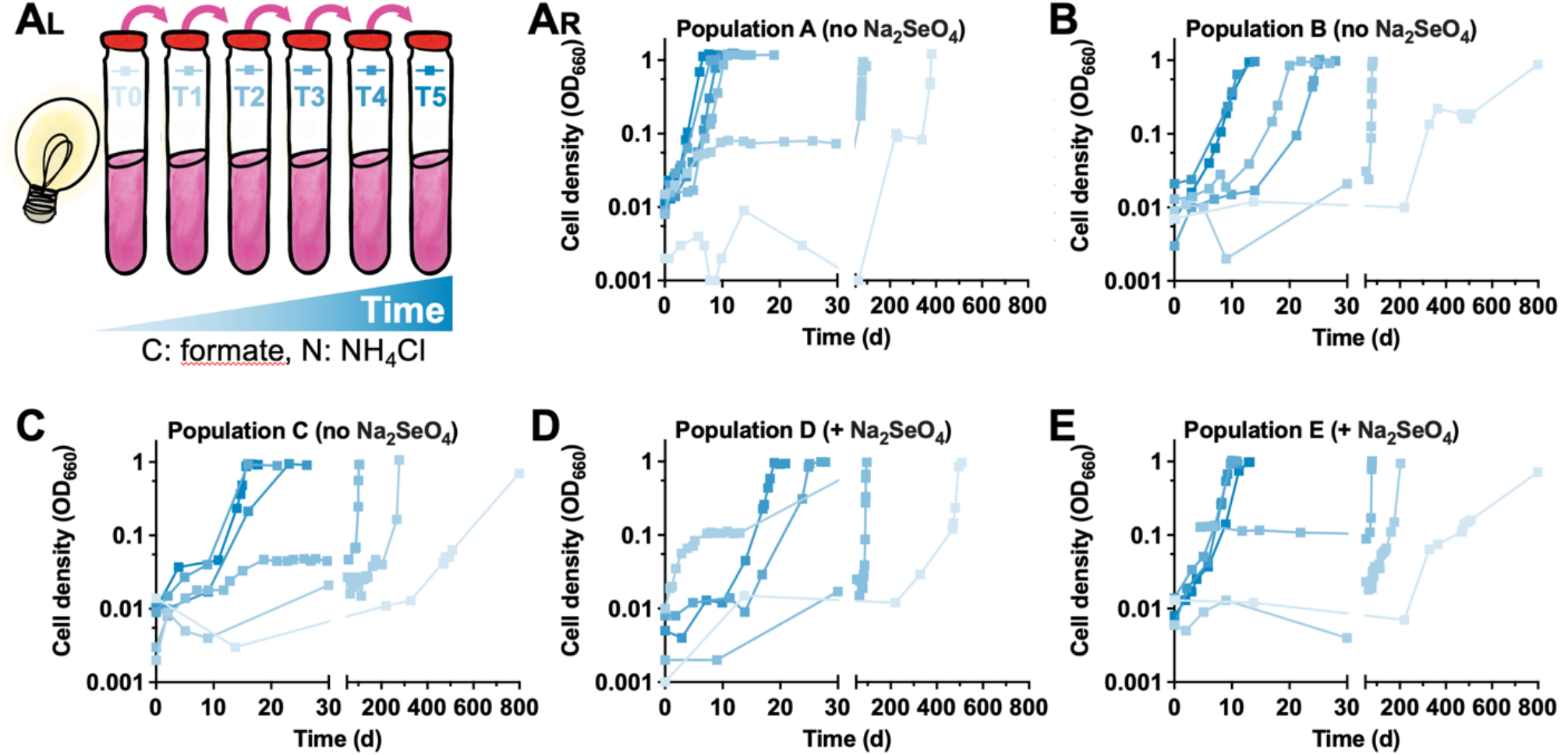
*R. palustris* NifA* can be evolved to use formate as the sole carbon source. **(A**_**L**_) *R. palustris* CGA4005 was serially transferred in a minimal medium with formate as the sole carbon source under constant illumination, with or without sodium selenate (Na_2_SeO_4_). Transfer (T) numbers 1-5 also received 5 mM thiosulfate. **(A**_**R**_ **– E)** Growth curves of populations A-E through serial transfers. Darker shading indicates later transfers.

Here we used adaptive laboratory evolution to evolve *R. palustris* isolates that use formate as the sole carbon source under anaerobic, phototrophic conditions. Hypothesis-driven gene deletions revealed that formate is assimilated via FDH and the CBB cycle. However, this pathway was not obviously connected to potentiating, actualizing, and refining mutations that, respectively, impacted photopigment production, flavin biosynthesis, and a regulator of unknown function. Thus, even when expansion of nutritional traits involves conventional pathways, the enabling mutations are not always intuitively connected. In this case, our results point to pigment production and chromatophore membranes as a possible CO_2_-concentrating mechanism.

## RESULTS

### Evolved formate-assimilating mutants contain common mutations

We explored the possibility of *R. palustris* formate assimilation using an ‘ancestor’ (CGA4005) that had a *nifA*\* mutation, which causes constitutive nitrogenase activity (N_2_ + 16 ATP + 8e^-^ + 10 H^+^ → 2NH_4_^+^ + 16 ADP + H_2_) (25, 26); we commonly use NifA* strains in coculture with fermentative *E. coli*, where formate accumulates (25). We incubated CGA4005 in anoxic defined medium with formate as the sole carbon source under constant illumination for phototrophic growth (Fig 1A). Formate oxidation by another Rhizobiales, *Sinorhizobium meliloti*, relies on a selenocysteine-containing FDH (21). Even though the *R. palustris* FDH was not predicted to have selenocysteine, we included selenate in three of the six cultures in case the prediction was wrong. Growth was detected in five cultures, starting 1-2 years after inoculation (Fig 1). Thus, selenate was neither sufficient nor required for formate utilization. Growth rates improved and lag times decreased through serial transfers (Fig 1).

To identify enriched mutations, we sequenced gDNA from two populations and five evolved isolates (one from each lineage; Fig. 1). Four genes were commonly mutated (Table 1): (i) *nifA*, encoding the transcriptional activator of nitrogenase, (ii) *ribB*, encoding 4-dihydroxy-2-butanone 4-phosphate (DHBP) synthase involved in flavin synthesis, (iii) *ppsR2*, an O_2_-sensitive repressor of light harvesting genes (27), and (iv) RPA0853 (OR798_04395), a predicted dual sensor histidine kinase-response regulator. Below we investigate the contribution of mutations in these genes to formate utilization.

**Table 1.** Mutations in formate-evolved isolates.

| Evolved Isolate | Position | Mutation | Annotation | Gene | Description |
| --- | --- | --- | --- | --- | --- |
| A | 5,211,791 | A→C | L563R (CTC→CGC) | *nifA* | transcriptional activator NifA |
| B | 5,211,791 | A→C | L563R (CTC→CGC) | *nifA* |  |
| C | 5,212,780:1 | +G | coding (699/1755 nt) E315* | *nifA* |  |
| D | 5,211,509 | Δ2,173 bp |  | *nifA* |  |
| E | 5,211,791 | A→C | L563R (CTC→CGC) | *nifA* |  |
| A | 944,428 | +C | coding (3458/3510 nt) +11 a.a. | *fbf*, OR798_RS04395 | PAS domain-containing hybrid sensor histidine kinase/response regulator |
| B | 946,303 | Δ7 bp | coding (1577-1583/3510 nt) D533* | *fbf*, OR798_RS04395 |  |
| C | 947,732 | A→T | W52R (TGG→AGG) | *fbf*, OR798_RS04395 |  |
| D | 945,373 | G→T | S838* (TCG→TAG) | *fbf*, OR798_RS04395 |  |
| E | 944,428 | +C | coding (3458/3510 nt) + 11 a.a. | *fbf*, OR798_RS04395 |  |
| A | 1,703,267 | G→A | G39E (GGG→GAG) | *ppsR2* | transcriptional regulator PpsR2 |
| B | 1,704,356 | T→A | I402N (ATT→AAT) | *ppsR2* |  |
| C | 1,704,361 | G→A | E404K (GAG→AAG) | *ppsR2* |  |
| D | 1,704,458 | A→G | K436R (AAA→AGA) | *ppsR2* |  |
| E | 1,703,267 | G→A | G39E (GGG→GAG) | *ppsR2* |  |
| A | 1,197,834 | C→T | R201C (CGC→TGC) | *ribB* | 3,4-dihydroxy-2-butanone-4-phosphate synthase |
| B | 1,198,239 | T→C | S336P (TCG→CCG) | *ribB* |  |
| C* | 1,198,114 | C→T | P294L (CCG→CTG) | *ribB* |  |
| C* | 1,198,227 | C→T | H332Y (CAT→TAT) | *ribB* |  |
| D | 1,198,252 | A→G | Y340C (TAC→TGC) | *ribB* |  |
| E | 1,197,834 | C→T | R201C (CGC→TGC) | *ribB* |  |

\* Evolved Isolate C has two populations of *ribB* mutations. P294L is at 74.4 frequency and H332Y is at 35.3 frequency.

### The *nifA*\* allele prevents formate assimilation

All evolved isolates had a mutation in *nifA*. The *nifA* mutations in isolates C and D respectively caused a frameshift and deleted most of the coding region (Table 1). These were loss-of-function mutations; isolates did not grow with N_2_ as the nitrogen source. However, the other three evolved isolates, which all had a NifA*^L563R^ mutation (Fig 2B), grew with N_2_ (Fig 2A).

**Fig 2.**
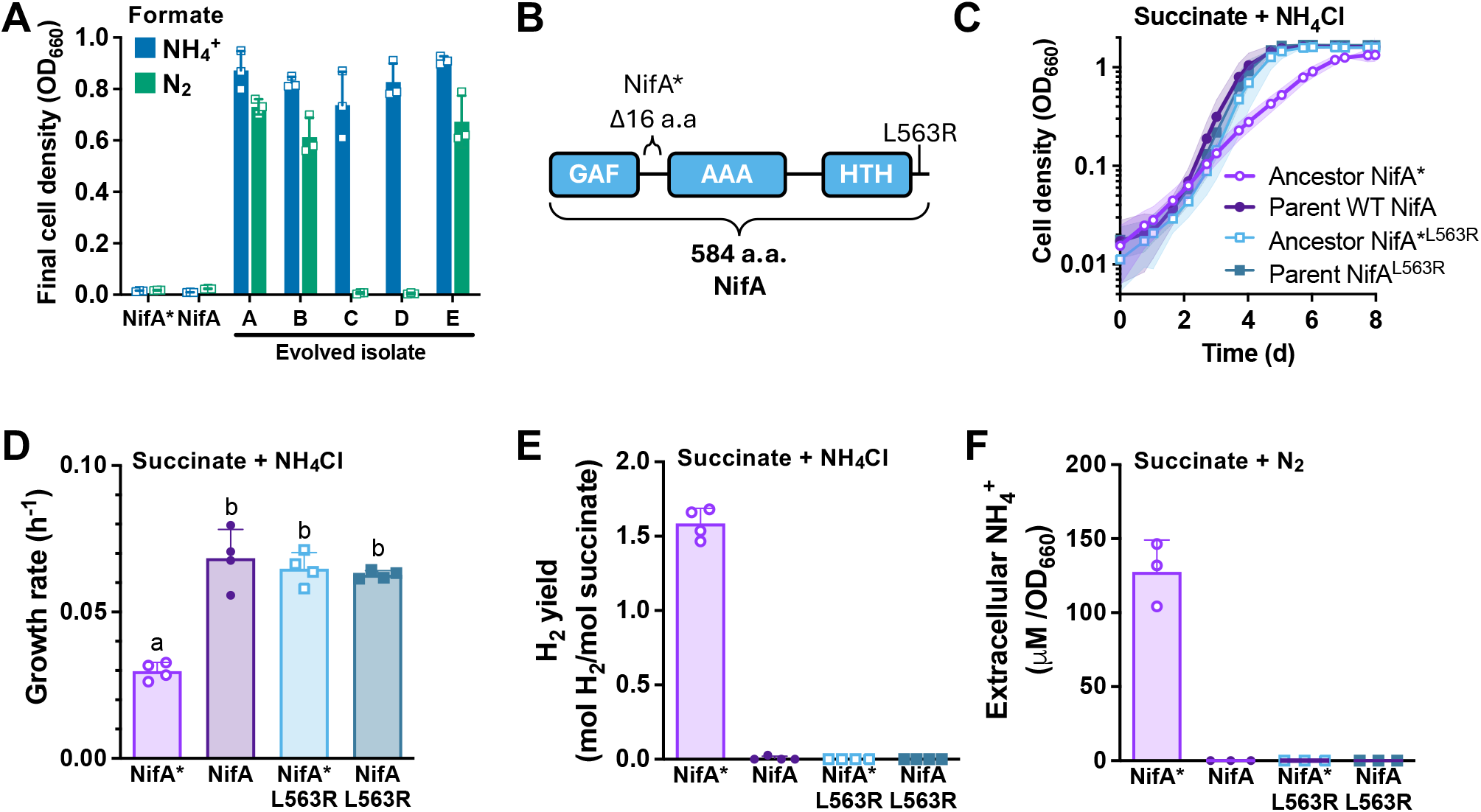
NifA mutations either eliminate nitrogenase activity or undo the constitutive NifA* phenotype and restore nitrogenase regulation. **(A)** Final cell densities with either NH_4_Cl or N_2_ as the sole nitrogen source and formate as the sole carbon source. **(B)** Location of the C-terminal NifA^L563R^ mutation in evolved isolates A, B, and E. The ancestor (CGA4005) has a 16 amino acid (a.a.) deletion that imparts constitutive nitrogenase activity (NifA*), resulting in slower growth and H_2_ production when cultured with NH_4_Cl, and NH_4_^+^ production when N_2_ is the sole nitrogen source. The parent (WT *nifA*; CGA4004) only expresses nitrogenase if N_2_ is the sole nitrogen source. **(C)** Growth curves with succinate as the sole carbon source and NH_4_Cl as the nitrogen source. The NifA^L563R^ allele was introduced into the parent (WT *nifA*, CGA4004) or the *nifA*\* ancestor (CGA4005). **(D, E)** Growth rates **(D)** and H_2_ yields **(E)** from the cultures in panel C. **(F)** NH_4_^+^ production with succinate as the carbon source and N_2_ as the nitrogen source. Error bars **(A, D-F)** or shading **(C)** = SD; n= 4. **(D)** Different letters indicate statistically different mean values (p < 0.05) as determined by one-way ANOVA with a Tukey multiple comparison post-test (Graphpad Prism v10).

Constitutive nitrogenase activity from the *nifA*\* allele causes slower growth accompanied by H_2_ production when NH_4_^+^ is the nitrogen source (26) and NH_4_^+^ excretion when N_2_ is the nitrogen source (25). To determine whether the L563R mutation affected these phenotypes, we moved the mutation into the NifA* ancestor, creating NifA*^L563R^ and into the ‘parent’ strain (WT *nifA*, CGA4004), creating NifA^L563R^. Each mutant exhibited growth trends with succinate and NH_4_Cl that resembled those of the WT NifA parent rather than the NifA* ancestor (Fig 2C, D). This observation suggested that the NifA*^L563R^ mutation bypassed nitrogenase misregulation. Indeed, neither mutant produced H_2_ when growth with NH_4_Cl (Fig 2E) nor did they excrete NH_4_^+^ when grown with N_2_ (Fig 2F).

Although nitrogenase is costly, we doubted that energetic cost selected for the NifA*^L563R^ allele because no *nifA* mutations were observed during separate long-term NifA* cultures with other carbon sources that lasted hundreds of generations (28, 29). We thus suspected that the NifA*^L563R^ mutation overcame a barrier to formate assimilation.

### Formate is assimilated via FDH and the CBB cycle

NifA* mutants have low CBB cycle gene expression and activity (26, 30). We thus hypothesized that formate is assimilated via CO_2_ using the CBB cycle. To test this hypothesis, we deleted the essential CBB cycle gene encoding phosphoribulokinase (Δ*cbbP*::km^R^) (31) in evolved isolate B (EIB). Deletion of *cbbP* abolished EIB growth with formate and NH_4_Cl (Fig 3A). The CBB cycle is important for both CO_2_ assimilation and maintaining electron balance when organic substrates are provided (26). Therefore, we also tested growth under N_2_-fixing conditions wherein nitrogenase provides an alternative means for electron balance (31). EIB growth with N_2_ was also abolished by the Δ*cbbP* mutation (Fig 3B). With both nitrogen sources, EIB growth was restored when *cbbP* was expressed from a plasmid (complemented) but not when an empty vector was used (Fig 3A, B). Thus, the CBB cycle was required to assimilate formate via CO_2_, and the lack of Δ*cbbP* mutant growth was not due to an inability to balance electrons.

**Fig 3.**
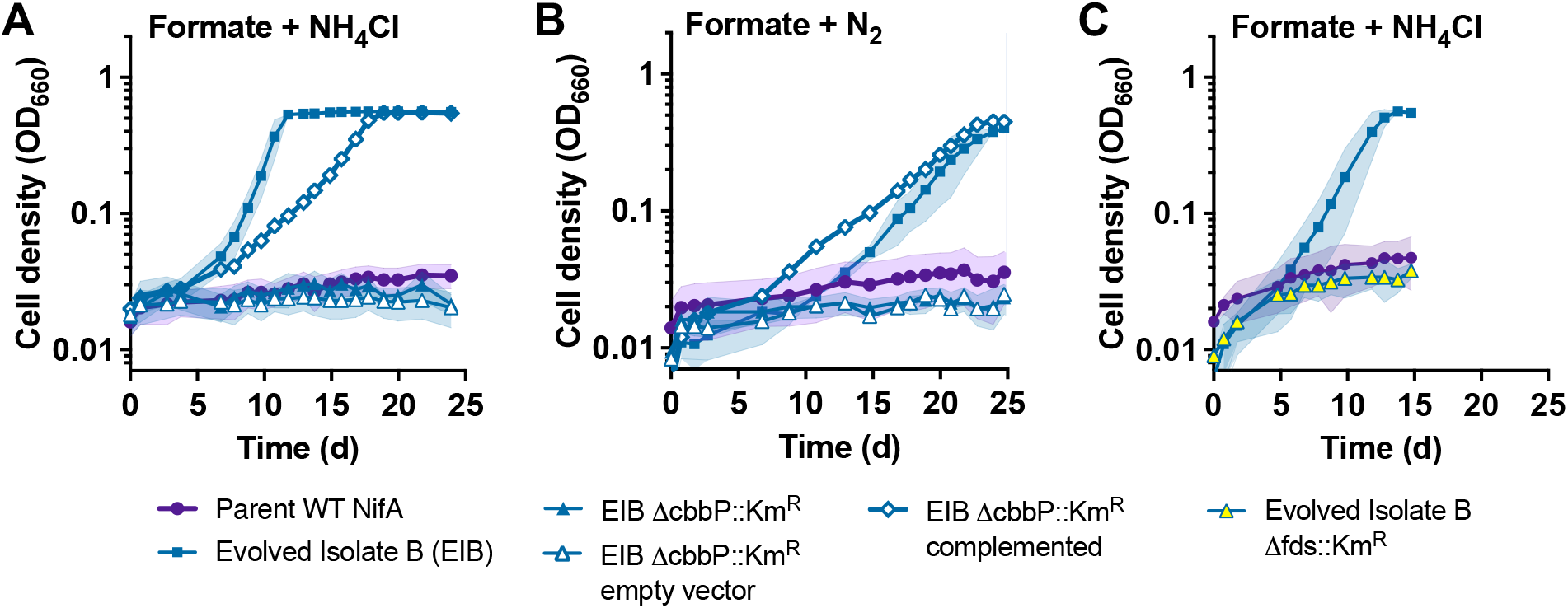
Formate is assimilated via FDH and the CBB cycle. **(A**-**C)** Growth with formate as the sole carbon source with or without the CBB cycle gene *cbbP* or the formate dehydrogenase genes *fdsGBA*. **(A)** Single representatives of triplicate EIB Δ*cbbP*::Km^R^ complemented cultures are shown due to variable lag times (0.1 OD_660_ reached between 12 - 17 h with NH_4_Cl or 15 - 30 h with N_2_ across all replicates). For all other growth curves, shading = SD; n = 3. Empty vector, pBBPgdh; complementation vector, pBBP*cbbP*; Parent WT NifA, CGA4004.

We hypothesized that *R. palustris* converts formate to CO_2_ using a cytoplasmic FDH (*fdsGBA*, RPA0732-4). Deleting *fdsGBA* in EIB abolished growth on formate (Fig 3C). We were unable to create a complementation vector despite multiple attempts. Still, our data strongly suggest that formate is assimilated via oxidation to CO_2_ by FDH and then CO_2_ fixation by the CBB cycle.

### NifA mutations were potentiating for formate assimilation

The lack of parent growth on formate, despite the WT *nifA* allele, indicates that *nifA* mutations alone are insufficient for formate assimilation. But the common occurrence of *nifA* mutations suggests that they set the stage for mutations that enabled formate assimilation. To test this hypothesis, we incubated the type strain (WT NifA; CGA0092), and two different CGA0092 Δ*nifA* strains (full vs partial deletion), with formate as the sole carbon source to enrich for formate-assimilating mutants. Both the WT *nifA* and Δ*nifA* alleles were insufficient for formate utilization (Fig 4). However, these backgrounds enriched for formate-utilizing mutants much faster (∼6 months; Fig 4). Two of the three evolved populations again had mutations in *ribB, ppsR2*, and RPA0853; *ribB* and/or *ppsR2* mutations swept to dominate early, whereas RPA0853 mutations appeared later (Table 2). The third population, originating from the partial Δ*nifA* strain, had a unique set of mutations involving *rpoD*, encoding the ‘housekeeping’ sigma factor, and *fixR2* (RPA4633, OR798_24065) encoded just upstream of *nifA* (Table 2). We leave the analysis of this unique mutant for a separate study, but we note that *rpoD* mutations were enriched in some long-term cocultures where formate levels were low (28). Our results suggest that the *nifA*\* allele in our original enrichments lowered the odds of enriching for formate-assimilating mutants, but the allele did not necessarily dictate the spectrum of mutations involved.

**Table 2.** Mutations in WT (F), *ΔnifA* full (G) and partial *ΔnifA* (H) populations.

| Population, transfer # | Position | Mutation | Annotation | Freq (%) | Gene | Description |
| --- | --- | --- | --- | --- | --- | --- |
| Wild-type (F), T0 | 1,703,446 | G→A | A99T (GCG→ACG) | 100 | *ppsR2* | transcriptional regulator of photosynthesis genes |
| T2 | 1,703,446 | G→A | A99T (GCG→ACG) | 100 | *ppsR2* |  |
|  | 1,198,233 | A→G | T334A (ACC→GCC) | 100 | *ribB* | riboflavin biosynthesis, DHBP synthase |
|  | 944,437 | Δ1 bp | coding (3449/3510 nt) L1156* | 58 | *fbf*, OR798_RS04395 | unknown function |
| full $\Delta nifA$ (G), T0 | 1,703,384 | Δ18 bp | coding (233-250/1395 nt) ΔI78-M81 | 100 | *ppsR2* | transcriptional regulator of photosynthesis genes |
|  | 1,197,935 | C→A | H234Q (CAC→CAA) | 100 | *ribB* | riboflavin biosynthesis, DHBP synthase |
| T3 | 1,703,384 | Δ18 bp | coding (233-250/1395 nt) ΔI78-M81 | 100 | *ppsR2* | transcriptional regulator of photosynthesis genes |
|  | 1,197,935 | C→A | H234Q (CAC→CAA) | 100 | *ribB* | riboflavin biosynthesis, DHBP synthase |
|  | 947,488 | A→G | L133P (CTG→CCG) | 15 | *fbf*, OR798_RS04395 | unknown function |
| partial $\Delta nifA$ (H), T0 | 1,407,234 | T→C | L395P (CTC→CCC) | 100 | *rpoD* | RNA polymerase sigma factor RpoD |
|  | 5,213,821 | T→C | E264E (GAA→GAG) | 100 | *fixR2*, OR798_RS24065 | SDR family oxidoreductase |
| T2 | 1,407,234 | T→C | L395P (CTC→CCC) | 100 | *rpoD* | RNA polymerase sigma factor RpoD |
|  | 5,213,821 | T→C | E264E (GAA→GAG) | 100 | *fixR2*, OR798_RS24065 | SDR family oxidoreductase |

**Fig 4.**
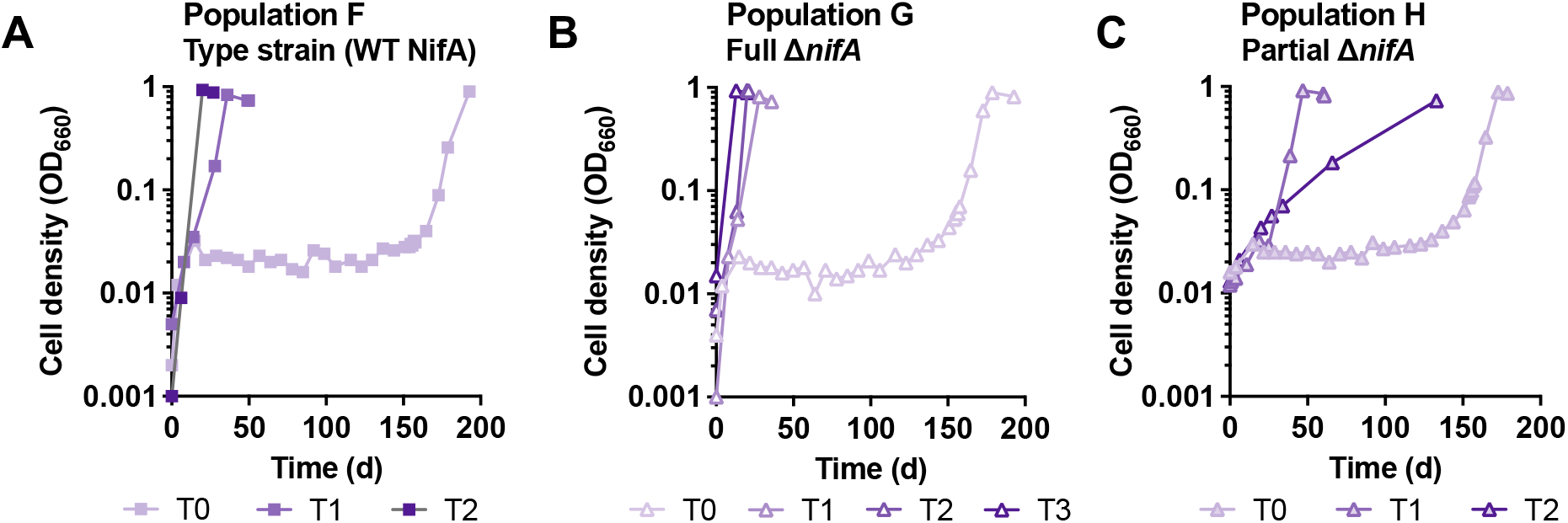
*R. palustris* can be evolved to use formate as the sole carbon source more quickly when the ancestor is not a NifA* strain. Type strain *R. palustris* CGA0092 (**A**; population F), a full Δ*nifA* strain (**B**; CGA4028; population G), and a partial Δ*nifA* strain (**C**; CGA757-2; population H), were serially transferred in sealed anaerobic test tubes in a minimal medium with formate as the sole carbon source and NH_4_Cl as the nitrogen source under constant illumination. Darker shading indicates later transfers (T).

### A RibBX mutation is necessary and sufficient for formate assimilation

As *nifA* mutations were insufficient for formate utilization (Fig 4) we turned our attention to the other common mutations. Each line had at least one mutation in *ribB*, encoding the flavin synthesis enzyme DHBP synthase. *R. palustris* DHBP synthase has a RibX domain (32) (Fig 5A). RibBX proteins are often misannotated as DHPB synthase/GTP cyclohydrolase II (RibBA). However, the RibX domain does not have GTP cyclohydrolase activity and likely regulates the DHBP synthase activity (32-34). Each *ribB* mutation caused a single amino acid substitution in the RibX domain (Table 1).

**Fig 5.**
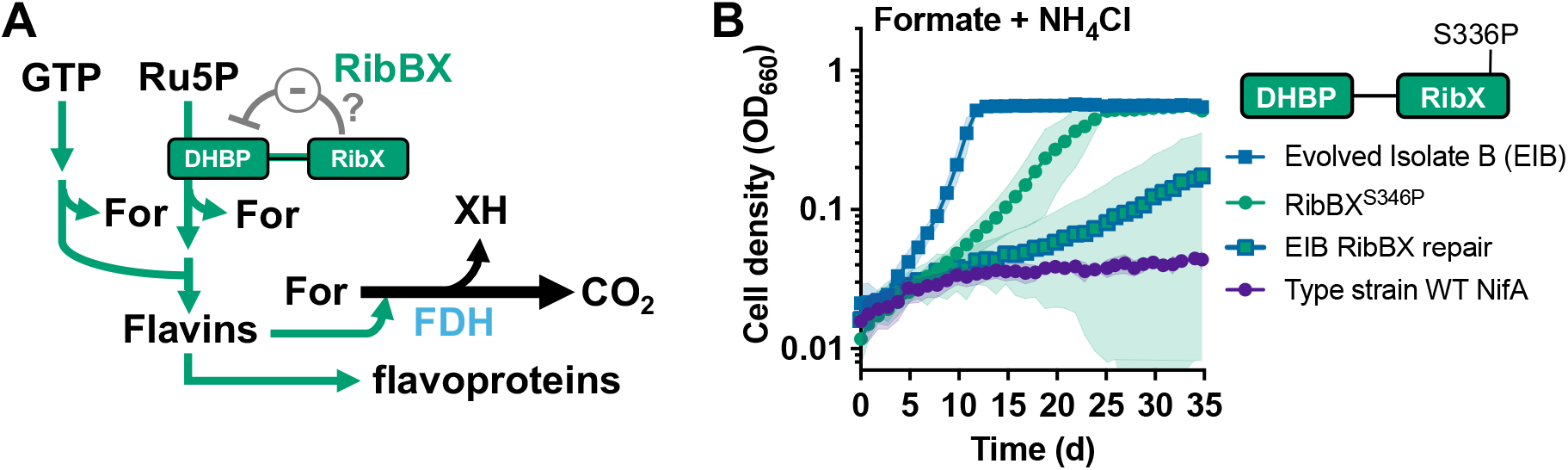
RibBX^S346P^ is sufficient for growth with formate as the sole carbon source. RibBX is involved in flavin biosynthesis (e.g., riboflavin, FMN, and FAD), and produces formate as a byproduct. FDH is a flavoprotein. The RibX domain is hypothesized to regulate 3,4-dihydroxy-2-butanone-4-phosphate (DHBP) synthase activity. For, formate; Ru5P, ribulose 5-phosphate; XH, reduced electron carrier. **(B)** Growth curves showing the effect of the RiBX^S346P^ mutation when introduced into the type strain (WT *nifA*, CGA0092) or when repaired in evolved isolate B (EIB). Shading = SD; n=4. Growth curves for the type strain and evolved isolate B are the same as in Fig 7. Biological replicates for the EIB RibX repair are shown in Fig S2.

Introduction of RiBX^S346P^ from EIB into the type strain (CGA0092, WT *nifA* allele), was sufficient for growth with formate as the sole carbon source, although the growth rate was lower than that of EIB (Fig 5B). RiBX^S346P^ was thus an actualizing mutation. Repairing the RiBX^S346P^ mutation in EIB eliminated growth in one replicate, but the other three replicates grew after different lag times (Fig 5B standard deviation; Fig S2). The variable lag times could suggest enrichment of new *ribB* mutants, perhaps facilitated by potentiating EIB mutations, like those in *nifA* and *ppsR2* (see below).

We considered how *ribB* mutations might interact with FDH or the CBB cycle. Related bacteria can use formate as an electron donor for autotrophic growth when supplemented with CO_2_ (21, 22). Thus, to test whether the RiBX^S346P^ mutation favors formate-oxidation as opposed to CO_2_ fixation, we cultured the parent (WT *nifA* allele), EIB, and the RiBX^S346P^ mutant under autotrophic conditions (NaHCO_3_ as the CO_2_ carbon source) with either formate or thiosulfate as electron donors (Fig S3). The parent used formate as an electron donor for CO_2_ fixation, but the EIB and RiBX^S346P^ mutants respectively grew 6.9- and 4.7-fold faster (Fig 6A, C). When thiosulfate was the electron donor, the EIB and the RiBX^S346P^ mutant showed respectively worse or similar growth compared to the parent (Fig 6B, C). These results indicate that RiBX^S346P^ facilitates formate oxidation but not CO_2_ fixation.

**Fig 6.**
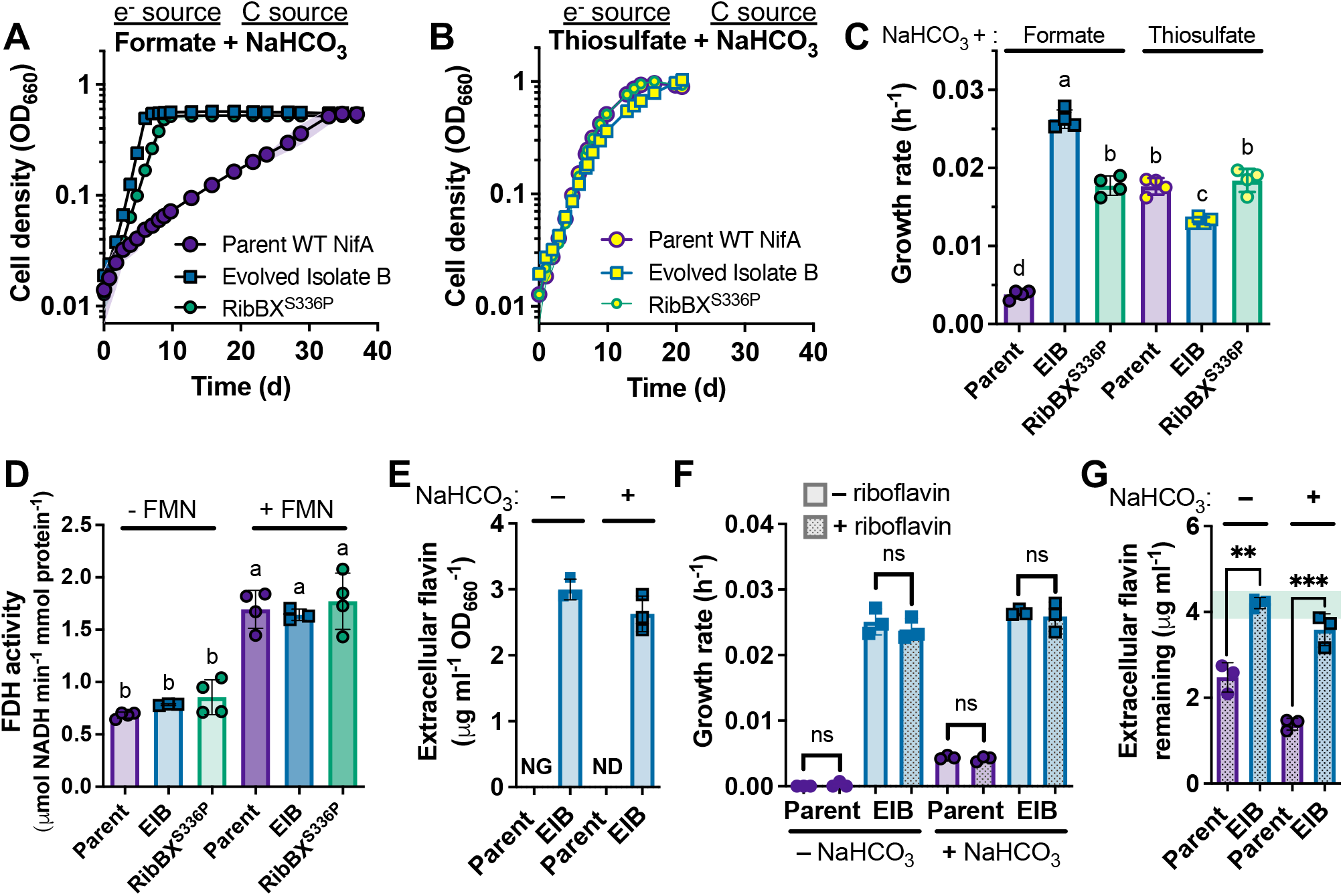
The RiBX^S346P^ mutation improves formate utilization during autotrophic growth. **(A**, **B)** Growth curves comparing the effect of all EIB mutations or just the RibBX^S336P^ on autotrophic growth when either 20 mM formate **(A)** or 20 mM thiosulfate as the electron source and NaHCO_3_ as the main carbon source (CO_2_ source). All mutations were made in the parent (WT *nifA*; CGA4004). Shading = SD; n=4. **(C)** Exponential growth rates for the cultures shown in panels A and B. **(D)** FDH activity assays in crude cell extracts. Values were determined by linear regression of A_340_ over time, subtracting background activity from extracts without formate. **(E)** Supernatant flavin concentrations from fully-grown cultures grown with formate with or without 20 mM NaHCO_3_; NG, no growth; ND, below the detection limit. **(F)** Exponential growth rates for cultures grown with formate +/- 20 mM NaHCO_3_ and +/- 4 µg/ml riboflavin. **(G)** Remaining riboflavin in supernatants from the cultures in panel F that were supplemented with riboflavin. **(C-G)** Error bars, SD; n=3 or 4. (**C, D)** Different letters indicate statistically different mean values (p < 0.05) as determined by one-way ANOVA with a Tukey multiple comparison post-test. **(F, G)** Unpaired t tests; ns, not significant, **, p < 0.01; ***, p < 0.001. All statistical tests were made using Graphpad Prism v10.

We hypothesized that improved formate oxidation could be attributed to increased FDH activity, which uses the flavin cofactor FMN (35) (Fig 5A). However, when we assayed FDH in crude cell extracts, the activity was similar between the WT, RibB^S346P^, and EIB strains (Fig 6D). Free FMN or FAD can stimulate FDH activity (36). When we added FMN to the FDH assays, activity increased in each extract by 2.1 – 2.5-fold (Fig 6D). Thus, although FDH activity was similar in each extract in vitro, RibB^S346P^ could have stimulated FDH activity by increasing in vivo flavin availability.

In *S. oneidensis*, the RibX domain inhibits DHBP synthase activity (Fig 5A) and removal of the RibX domain increases flavin externalization (32). Similarly, we detected flavins in supernatants of formate-grown EIB cultures (Fig 6E) whereas no flavins were detected in parent cultures grown with formate plus NaHCO_3_. Thus, the *R. palustris* RibB^S346P^ mutation likely increased DHBP synthase activity. Based on this result, we hypothesized that access to flavins would allow the parent to grow with formate. However, supplementing parent cultures with riboflavin did not enable growth on formate without NaHCO_3_, nor did it affect the parent growth rate with formate plus NaHCO_3_ (Fig 6F). We assume that *R. palustris* can take up riboflavin because stationary phase concentrations were below the initial 4 µg/ml (Fig 6G) and because concentrations were stable in uninoculated controls. However, the non-growing *R. palustris* parent removed nearly as much riboflavin as a fully grown parent culture (Fig 6G), which could suggest biodegradation rather than utilization as a cofactor. EIB stationary phase flavin levels were similar to the amount initially added, possibly suggesting a steady state between flavin externalization and uptake.

We also considered that the RibX domain could repress DHBP synthase in response to formate via feedback inhibition; formate is a product of DHPB synthase (Fig 5A). If so, using millimolar concentrations of formate as a carbon source could inhibit flavin synthesis and thereby growth. However, if true, we expect that a riboflavin supplement would have circumvented this inhibition, whereas riboflavin had no effect on parent nor EIB growth (Fig 6F).

### PpsR2 mutations facilitate formate utilization

Another commonly mutated gene was *ppsR2*, encoding an O_2_-responsive repressor of light-harvesting genes, such as those for bacteriochlorophyll (BChl), carotenoid synthesis, and light harvesting complexes in intracellular chromatophore membranes (37) (Fig 7). In some evolved populations, *ppsR2* mutations were enriched to 100% before *ribB* and/or RPA0853 mutations (Populations B and D at transfer 3 and F and G at transfer 0; Table S1). However, *ppsR2* mutations commonly emerge in *R. palustris* cultures, independent of formate assimilation (28, 29); the increased pigmentation (Fig S4) is likely advantageous during light limitation at high cell densities (27). We therefore questioned whether *ppsR2* mutations are important for formate utilization.

**Fig 7.**
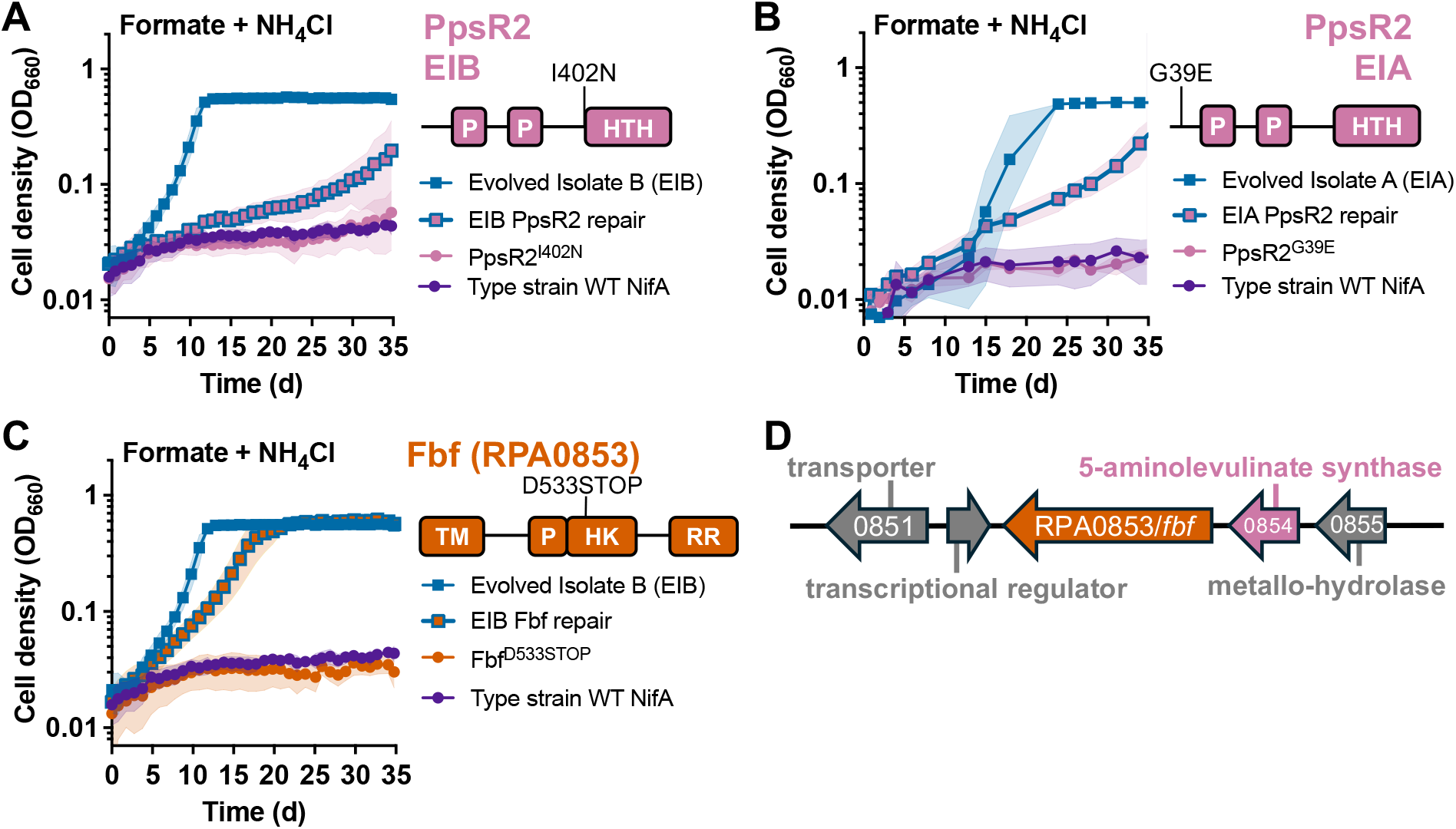
PpsR2 mutations and RPA0853^D533STOP^ mutations facilitate growth with formate as the sole carbon source. **(A, B)** Growth curves showing the effect of each *ppsR2* mutation when introduced into the type strain (WT *nifA*; CGA0092) or when repaired in either evolved isolate B **(A)** or evolved isolate A **(B). (C)** Growth curves showing the effect of the RPA0853^D533STOP^ mutation when introduced into the type strain (WT *nifA*, CGA0092), or when repaired in evolved isolate B (EIB). **(A-C)** Shading = SD; n=4. Growth curves for EIB and the parent strain in panels A and C are the same for as in Fig 5. **(D)** RPA0853 (*fbf*) is encoded downstream of a gene that potentially encodes 5-aminolevulinate synthase, involved in BChl synthesis.

When we introduced PpsR2^I402N^ from EIB into the parent, the mutant did not grow with formate as the sole carbon source (Fig 7A). However, repairing the mutation in EIB negatively impacted growth (Fig 7A). The same tests with the mutation from EIA gave similar results (Fig 7B). With the caveat that we did not test all mutations, *ppsR2* mutations alone are insufficient but facilitate formate assimilation and likely serve as a potentiating mutation, given the early emergence.

### RPA0853 mutations facilitate formate assimilation

RPA0853 is a predicted dual histidine kinase response regulator and Na+/solute transporter (Fig 7C). RPA0853 mutations appeared later than those in *ribB* and *ppsR2* (Table 2). Introducing RPA0853^D533STOP^ into the parent was insufficient for growth with formate as the sole carbon source (Fig 7C). However, repairing the mutation in EIB negatively impacted growth (Fig 7C). Thus, the mutation facilitates growth with formate, likely as a refining mutation given the late emergence. We propose naming RPA0853 *fbf* for formate beneficial factor.

### Possible role of PpsR2 and Fbf in Rubisco priming

We considered how PpsR2 and Fbf might facilitate formate utilization. To assimilate formate via the CBB cycle, CO_2_ must first accumulate to levels that can be captured by Rubisco. Increasing CO_2_ availability would explain the upward-curving growth trends we often observed, rather than the expected linear trend for a constant exponential growth rate on a semi-log plot (Fig 1, 3-5, 7).

To gauge how CO_2_ availability might affect growth trends, we simulated cultures using a Monod model that included our measured growth yields and rates and published FDH and Rubisco half-saturation constants (Km) (36, 38, 39) (Fig 8A). We assumed that reductant (XH) was initially available and could accumulate without consequence. We then varied the ratio between the formate oxidation and CO_2_ fixation rates. Our simulations suggest that growth requires the formate oxidation rate to exceed the CO_2_ fixation rate, thereby allowing CO_2_ to accumulate to levels that eventually drive Rubisco (Fig 8B); this is consistent with our findings that the RibB^S346P^ mutation improves formate oxidation (Fig 6). However, even a 1.7-fold higher formate oxidation rate (estimated to reflect EIB cultures) could not explain observed growth trends (Fig 8B, orange). If we included an initial 20 µM of CO_2_ (estimated HCO_3_^-^ carry-over from succinate-grown starter cultures (30)), then simulated growth occurred even when formate oxidation and CO_2_ fixation rates were equal, and a 1.3-fold higher formate oxidation rate explained observed trends (Fig 8C). Thus, a threshold amount of CO_2_ should be able to prime Rubisco to then sustain continuous formate oxidation.

**Fig 8.**
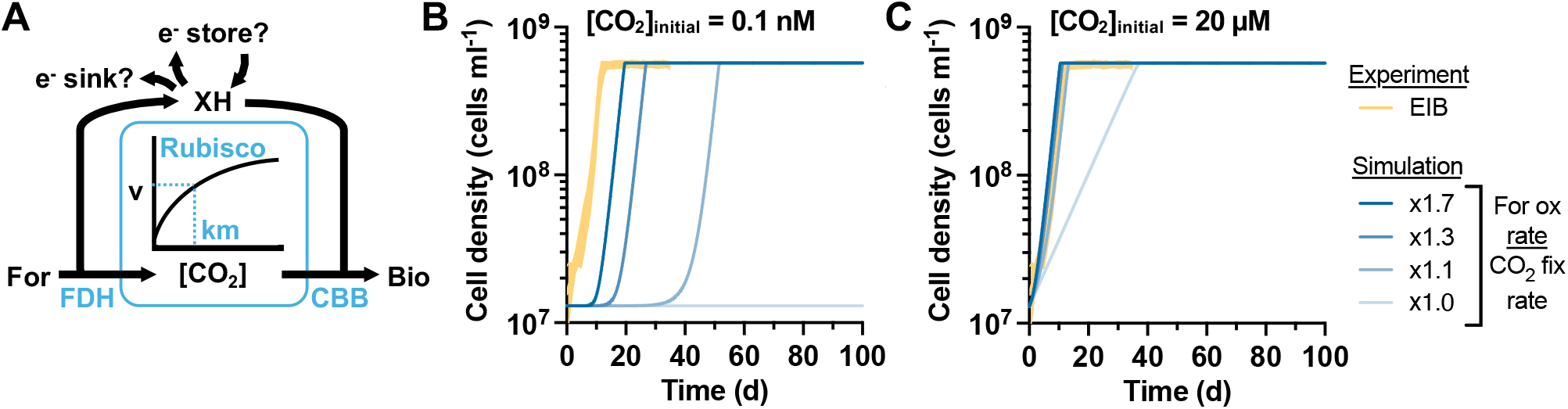
CO_2_ accumulation is necessary for utilizing formate as the sole carbon source. **(A)** Formate oxidation by FDH generates CO_2_ that must accumulate before significant Rubisco activity (v) can occur. Electrons liberated by formate oxidation are carried by electron carriers (XH). These electron carriers can donate electrons to the CBB cycle but might need to donate electrons elsewhere while CO_2_ accumulates. **(B, C)** Simulated cultures based on a Monod model where formate is the main carbon source with either 0.1 nM **(B)** or 20 µM **(C)** initial CO_2_; no growth occurred if the initial CO_2_ concentration was zero. The ratio of formate oxidation : CO_2_ fixation rate was varied by progressively lowering the growth yield on CO_2_, which we measured to be 1.7-times that of the growth yield on formate (Fig 6B vs Fig 3A). Orange, standard deviations from 10 independent EIB growth curves with formate as the sole carbon source (data from Fig 3, 5, and 7).

The simulated cultures assumed that electrons from formate oxidation could be stored or discarded while CO_2_ accumulated. H_2_ was reported as an electron intermediate by another *R. palustris* strain during growth with formate (20, 40), but our strains do not have formate hydrogen-lyase, nor did we detect H_2_ in the culture headspace. We therefore questioned (i) how electrons could be managed during CO_2_ accumulation and (ii) whether other mechanisms than FDH might generate CO_2_ without accompanying reductant. One possibility to address both challenges is elevated pigment production via *ppsR2* mutations (Fig S4). Synthesis of each BChl and carotenoid molecule would oxidize 10 and 8 net NAD(P)H equivalents, respectively (Fig 9A, XH). Synthesis of each BChl and carotenoid also, respectively, generates 18 and 4 CO_2_ from central metabolic precursors via redox-independent decarboxylases (Fig 9A). Although Fbf mutations did not noticeably alter pigmentation (Fig S4), Fbf might contribute to pigment synthesis, given its location immediately downstream of a possible 5-aminolevulinate synthase (RPA0854, Fig 7D), which generates 8 of the CO_2_ associated with BChl synthesis. RPA0854, shares 60% amino acid identity with another 5-aminolevulinate synthase, RPA1554, that is encoded in a BChl synthesis gene cluster.

**Fig 9.**
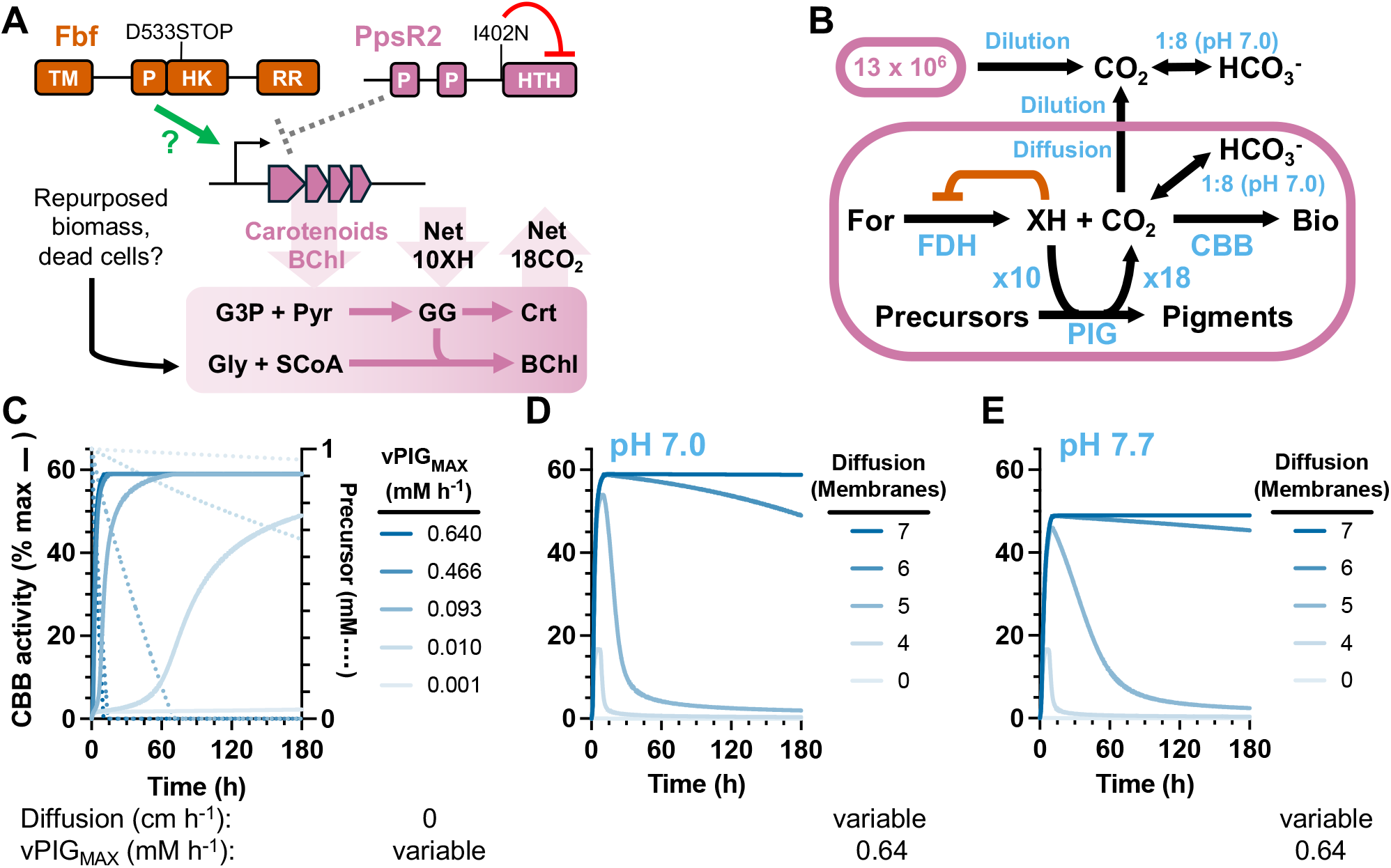
Simulations suggest that pigment synthesis and chromatophore membranes could form a CO_2_ concentrating mechanism. **(A)** Mutations in *ppsR2* relieve repression of pigment synthesis, which generally correlates with light harvesting complexes and chromatophore membranes. Fbf might also regulate the first step of BChl synthesis given its proximity to a pigment synthesis gene (Fig 7D). The synthesis of each BChl molecule uses 10 reducing equivalents (XH) and generates 18 CO_2_. The synthesis of each carotenoid uses 8 XH and generates 4 CO_2_ (not shown). **(B)** A single cell Michaelis-Menten model wherein pigment synthesis can both relieve inhibition of FDH by using XH and generate CO_2_ to prime Rubisco in the CBB cycle. CO_2_ is in equilibrium with HCO_3_^-^, catalyzed by carbonic anhydrase in the cell. CO_2_ can also diffuse out of the cell and spontaneously equilibrate with HCO_3_^-^ at a slower rate. We assumed that extracellular CO_2_ also came from 13 × 10^6^ cells in the inoculum, accounting for the dilution effect going from the volume of the cytoplasm to the volume of the medium. **(C-E)** Effect of the maximum pigment synthesis rate **(C)** the membrane permeability coefficient for CO_2_ based on the number of chromatophore membranes **(D)**, and a pH of 7.7 (∼1 CO_2_ : 40 HCO_3_^-^) **(E)** on the CBB cycle rate, using the specified parameter values below each graph. The permeability coefficients used for each number of membranes is in Table S4. **(C)** Precursor levels are also shown as dotted lines. **(D, E)** Precursors were exhausted in under 15 h for all conditions.

To explore the feasibility of pigment synthesis facilitating formate assimilation, we constructed a single-cell Michaelis-Menten model (Fig 9B). The model describes CO_2_ production from FDH and pigment synthesis, inter-conversion of CO_2_ and HCO_3_^-^ (∼1 CO_2_ : 8 HCO_3_^-^ at pH 7), and diffusion of CO_2_ from the cell. We also assumed that other cells in the inoculum contributed to extracellular CO_2_. We simulated 180 h to approximate the generation time for the parent strain with formate and NaHCO_3_. We initially set CO_2_ diffusion to zero. An initial precursor availability for pigment synthesis was necessary for CBB activity (Fig S5A). We set the precursor concentration to 1 mM for further simulations. A minimum pigment synthesis velocity (vPig_MAX_) of > 0.01 mM h^-1^ was also necessary for CBB activity (Fig 9C). Higher pigment synthesis velocities, based on estimates for the parent or the EIB strain with or without NaHCO_3_, achieved high CBB activity within 30 h (Fig 9C). CBB activity was maintained after precursors were exhausted (Fig 9C, dotted lines), suggesting that once Rubisco is primed, its activity can be sustained by CO_2_ and reductant from FDH.

Introducing CO_2_ diffusion using permeability coefficients for cholesterol-rich membranes (360 cm h^-1^) and carboxysomes (36 cm h^-1^) (23, 41) completely prevented CBB activity (Fig 9D). However, between the mid-cell and the cell envelope, *R. palustris* can have up to 16 layers of intracellular chromatophore membranes (42, 43), similar to thylakoid membranes that potentially limit CO_2_ diffusion in algal pyrenoids that house Rubisco condensates (44). Intracellular membrane surface area also correlates with the pigment levels (43, 45). Assuming each membrane is composed of phospholipids with 18-carbon fatty acids (46), and that each membrane decreases the permeability coefficient by a factor of 0.037 (47), our simulations suggest that 5 – 7 membranes could be sufficient to trap CO_2_ and prime Rubisco (Fig 9D).

pH can also play a CO_2_-concentrating role by determining the equilibrium between CO_2_ and HCO_3_^-^, the latter of which has a low membrane permeability. An alkaline pH in the chloroplast stroma traps HCO_3_^-^ around algal pyrenoids (24, 44), which are comparatively acidic due to proton release by Rubisco, thus favoring local conversion of HCO_3_^-^ to CO_2_ (48). We posit that the decarboxylases used in pigment synthesis could create alkaline conditions by consuming protons, analogous to decarboxylases that offset severe acid stress in *E. coli* (49). When we simulated a pH of ∼7.7 (1 CO_2_ : 40 HCO_3_^-^), the maximum CBB rate was lower, but the decline in CBB activity over time was less pronounced when ≥ 5 membranes were simulated (Fig. 9E). These results were not affected by CO_2_ generated from a population of cells due to the large dilution effect when CO_2_ escapes to the medium (Fig S5B). Together, these simulations suggest that a CO_2_-concentrating mechanism is required to grow on formate as the sole carbon source and that this requirement could be met by attributes associated with increased pigment synthesis such as: CO_2_ production, (iii) diffusion-limiting intracellular membranes, and (iv) HCO_3_^-^-trapping alkaline conditions.

## DISCUSSION

Here we used adaptive laboratory evolution to enrich for *R. palustris* mutants that (i) utilized formate as the sole carbon source (Fig 5) and (ii) exhibited improved autotrophic growth with formate as the electron donor (Fig 6). Formate was first oxidized by FDH to CO_2_, followed by assimilation via the CBB cycle, but the responsible mutations were not directly associated with genes for either pathway. Based on the time of mutation detection and the imparted phenotypes, the enriched mutations fit a similar sequence of potentiation (*nifA* and/or *ppsR2*), actualization (*ribB*), and refinement (*fbf*), as described for evolved citrate utilization in *E. coli* (7). Progressively increasing growth rates during enrichment, likely due to increasing CO_2_ availability for Rubisco, prevent us from confidently assessing whether potentiating, and possibly actualizing, mutations were enriched from a pre-existing subpopulation or if they arose during incubation (Fig S6). We favor the former. Although *R. palustris* has exceptional longevity under constant illumination (50), chromosome replication, and thus mutation generation, would be low without growth.

Our data suggest that the above mutations addressed several challenges such as: (i) ensuring the CBB cycle is active to assimilate CO_2_ (*nifA*) (ii) ensuring that formate oxidation meets CBB cycle demands for CO_2_ and/or reductant (*ribB*), (iii) providing an electron sink for formate oxidation when CO_2_ levels are insufficient to support CBB activity (*ppsR2* and possibly *fbf*), and (iv) providing an auxiliary source of CO_2_ to prime Rubisco (*ppsR2* and possibly *fbf*). The need for multiple parallel mutations indicates that the genetic landscape leading to formate assimilation was complex. Our observations demonstrate that novel nutritional traits can arise through unintuitive means, even when conventional pathways are involved. Understanding these connections should lead to more meaningful inferences for how novel nutritional traits evolve.

Early *nifA* mutations were likely potentiating when the ancestor had a *nifA*\* allele (Fig 2). The *nifA*\* allele causes constitutive nitrogenase activity and low CBB cycle activity (30). Thus, the ancestral *nifA*\* strain had an unanticipated regulatory barrier to formate assimilation via CO_2_. NifA^L563R^ is the first mutation known to bypass the *nifA*\* allele. Despite the slower growth rate of NifA* strains, the *nifA*\* allele was previously found to be stable through hundreds of generations in both monocultures and in cocultures with *E. coli* (28, 29). The NifA*^L563R^ mutation likely required the strong selective pressure exerted when the CBB cycle is required for growth on formate. It is currently unclear whether the mutation restores nitrogenase regulation or if it simply dampens NifA activity.

A mutation in the RibX domain of *ribB* was the only mutation we tested that was sufficient for growth on formate as the sole carbon source. The mutation affected formate oxidation, likely by increasing flavin availability for FDH (Fig 6). However, adding riboflavin did not improve growth with formate for any strain we tested. Thus, it is possible that the RiBX^S346P^ mutation affects formate utilization in ways that are independent of flavins.

Whereas the regulons of PpsR2 and Fbf deserve future scrutiny, we are intrigued by their connection to pigment synthesis, which could serve as a CO_2_-concentrating mechanism by (i) generating local CO_2_ (Fig 10), (ii) limiting CO_2_ diffusion via physical barriers of intracellular membranes, and (iii) creating local alkaline conditions that help trap CO_2_ as HCO_3_^-^. These features resemble algal pyrenoids, where Rubisco is concentrated as condensates in an acidic microenvironment (48) surrounded by starch, thylakoid membranes, and an alkaline stroma, all of which can help retain CO_2_ (24, 44). The similarities raise questions about whether a spatial arrangement of microenvironments also exists in *R. palustris* to facilitate CO_2_ fixation (Fig 10). Does Rubisco form condensates? Is Rubisco situated with carbonic anhydrase similar to the CO_2_-concentrating feature of carboxysomes (23)? Does *R. palustris* use inorganic carbon transporters and where are they situated (24, 44)? How does the dense array of light-harvesting proteins in chromatophore membranes (51) affect permeability?

**Fig 10.**
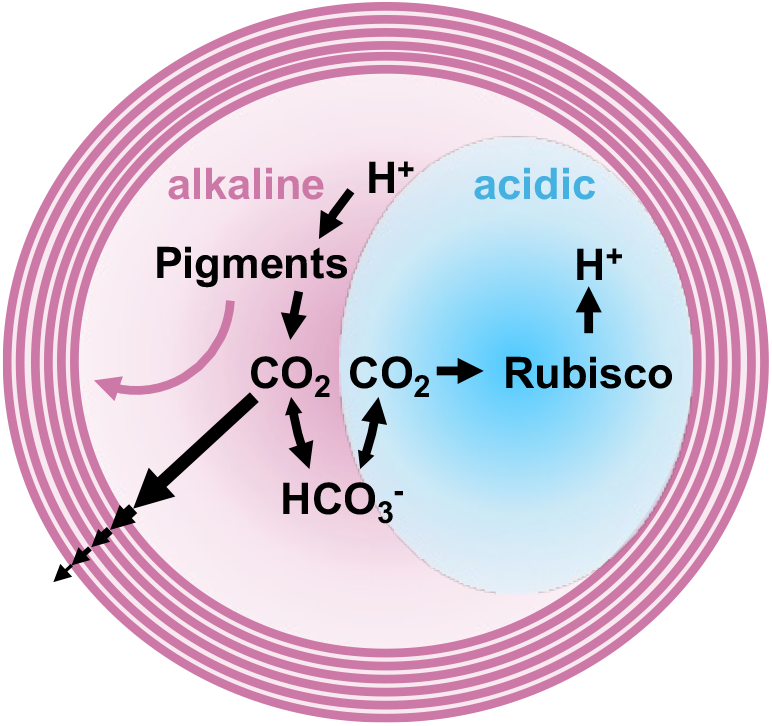
A hypothetical spatial coordination of pigment synthesis and Rubisco could facilitate CO_2_ concentration. Hypothetical axial view of an *R. palustris* cell, where pigment synthesis generates light-harvesting BChl and carotenoids for chromatophore membranes (pink arrow leading to pink membrane stacks). Multiple membrane stacks could serve to limit CO_2_ diffusion from the cell (decreasing arrow widths). Pigment synthesis also generates CO_2_ that could prime Rubisco and an alkaline microenvironment that can trap CO_2_ as HCO_3_^-^. Rubisco creates an acidic microenvironment while fixing CO_2_, helping convert local HCO_3_^-^ to CO_2_.

Proton-consuming decarboxylases associated with pigment synthesis are intriguing on their own as a rudimentary CO_2_-concentrating mechanism. Perhaps this feature explains why α-proteobacteria use the decarboxylating 5-aminolevulinate synthase as the first step to make BChl and other tetrapyrroles (e.g., heme), whereas other bacteria use a C5-pathway that does not liberate CO_2_ (52). Pigment-associated decarboxylases could be broadly beneficial to autotrophs. For example, pigment synthesis could contribute to stroma’s alkaline pH around the pyrenoid. In chemoautotrophs, the same decarboxylase reactions used in pigment synthesis would instead participate in the production heme and quinones used for respiration. Based on these features, one can speculate that a CO_2_-concentrating role associated with the synthesis of tetrapyrroles (e.g., BChl and heme) and isoprenoids (e.g., carotenoids and quinones) could have evolved alongside the energy-transforming roles of the final products.

## METHODS

### Strains and growth conditions

Strains, plasmids, and primers are in Table S2. All strains were derived from type strain CGA0092 (19). The ‘ancestor’ NifA* strain was CGA4005. The ‘parent’ strain (WT *nifA* allele) was CGA4004. CGA4004 and CGA4005 have a Δ*uppE* mutation, limiting biofilm formation (53), and a Δ*hupS* mutation, preventing H_2_ oxidation (25). CGA4005 has a 48-nucleotide deletion in *nifA* resulting in constitutive nitrogenase activity (26).

Unless indicated otherwise, *R. palustris* was cultured in 10 ml of defined photosynthetic medium (PM) (54, 55) in 27-ml anaerobic test tubes. For N_2_-fixing conditions, *R. palustris* was grown in PM without [NH_4_]_2_SO_4_ (N_2_-fixing media; NFM). Media were made anoxic by bubbling with Ar (PM) or N_2_ (NFM) and then sealing with a rubber stopper and an aluminum crimp before autoclaving. Starter cultures were inoculated from single colonies grown on PM agar with 10 mM disodium succinate. Test cultures received a 1% inoculum of stationary phase starter culture. All cultures were supplemented as indicated with either 10 mM succinate, 20 mM or 40 mM sodium formate, 5 or 20 mM sodium thiosulfate, and 20 mM NaHCO_3_ (56). Cultures were incubated upright without shaking (PM) or horizontally with shaking at 150 rpm (NFM) at 30°C in front of a 45 W halogen bulb (430 lumens). For cultures supplemented with riboflavin (4 µg/ml) cultures were incubated upright without shaking in front of a bank of eight infrared LED bulbs (850 nm, Univivi) to avoid photodegradation (57). Kanamycin (100 µg ml^-1^) or gentamycin (100 µg ml^-1^) were added where appropriate.

### Adaptive laboratory evolution

*R. palustris* CGA4005 starter cultures were inoculated in PM with formate as the sole carbon source, with or without 20 μM sodium selenate (Fig 1A). Stationary phase cultures were then serially transferred to fresh medium that included 5 mM sodium thiosulfate. Cultures from transfer 3 were plated on PM agar with succinate. One colony from each line was randomly selected, confirmed to use formate as the sole carbon source, and then genome sequenced. Two enrichments (populations B and D) were also selected for metagenome sequencing after transfer 3. Evolved cultures derived from CGA0092 and Δ*nifA* strains were serially transferred as above, without selenate. Populations were sequenced after three transfers.

### Genome sequencing

Genomic DNA (gDNA) was extracted from 1 ml of culture using a Qiagen DNeasy Blood and Tissue Kit. Lysis was facilitated using proteinase K (50 μg ml^-1^; 56°C for 10 min) followed by treatment with RNaseA (4 μl, Promega; 2 min). For evolved CGA4005 isolates and populations, DNA fragment libraries were prepared and sequenced by the IU Center for Genomics and Bioinformatics as described (29). For evolved CGA0092 and Δ*nifA* populations, libraries were prepared and sequenced on an Illumina NextSeq 2000 (2 × 151-bp reads) at SeqCenter (seqcenter.com) as described (19). Paired-end reads were pre-processed using cutadapt 3.4 (58). Mutations were identified using breseq v. 0.32.0 (59) using a concatenation of the CGA009 genome and plasmid (accession BX571963, BX571964) (29). Raw reads are in the NCBI Sequence Read Archive under BioProject accession number PRJNA1242330 (www.ncbi.nlm.nih.gov/bioproject).

### Construction of *R. palustris* mutants

Mutations from evolved isolates were moved into the ancestor (CGA4005), parent (CGA4004), or type strain (CGA0092), or repaired in evolved isolates by amplifying ∼0.5 kb flanking each side of a mutation using the designated primers (Table S2). Gene deletion mutants were made by amplifying 1 kb flanking the gene and then combining the products using overlap extension PCR as described (31). In each case, PCR products were then incorporated into linearized suicide vector pJQ200SK (60, 61) (along with a kanamycin resistance cassette for Δ*fdsGBA*::Km^R^) by Gibson assembly (New England Biolabs) or by restriction digest and ligation, maintained in *E. coli* NEB10β, and introduced into *R. palustris* by electroporation (62). Homologous recombinants were isolated by selective plating with gentamycin. Second recombinants were selected using 10% sucrose for counter selection and verified by PCR screening and Sanger sequencing.

### Analytical procedures

Cell densities were assayed in culture test tubes via optical density at 660 nm (OD_660_) using a Genesys 20 visible spectrophotometer (Thermo-Fisher). Specific growth rates were calculated using values between 0.1 and 1 OD_660_, where the relationship between OD and cell density is linear. H_2_ was quantified using a Shimadzu gas chromatograph as described (63). NH_4_^+^ was quantified using an indophenol colorimetric assay (25). Flavins were quantified in stationary phase supernatants using a Biotek Synergy H1 (excitation, 440 nm; fluorescence, 525 nm) (32). Riboflavin was used for the standard curve.

### FDH assay

Strains were grown in PM with 20 mM sodium formate and 20 mM NaHCO_3_ until late exponential phase (0.3-0.4 OD_660_). Cultures were pelleted by centrifugation at 4°C and then washed and resuspended in 1 ml of ice-cold 100 mM Tris/HCl, pH 7.5 + 10 mM MgCl_2_. Cells were lysed via three rounds of sonication with a Branson Sonifier (amplitude 35; 10 s bursts). Cells were chilled on ice for 1 min between rounds. Lysates were clarified by centrifugation. Assays were performed in a 96-well plate at 30°C in 250 µl with 5 mM NAD^+^, 60 mM sodium formate, and 10 µl of cell extract, with or without 1 µm FMN. A_340_ was converted to NADH (mM) using a standard curve.

### Simulations

Formate-assimilating cultures were simulated using a Monod model (Fig 10) and single-cell metabolism was simulated using a Michaelis-Menten model (Fig 11) using RStudio (64). Equations and default parameters are in the Supplementary materials. Some values were found via BRENDA (39), BioNumbers (65), or Biocyc (66). RStudio models are at https://github.com/McKinlab/Coculture-Mutualism.

## ACKNOWLEDGEMENTS

We are grateful to Ankur Dalia, Dan Kearns, Jay Lennon, Julia van Kessel, and McKinlay lab members for useful discussions. We also thank Caroline Harwood, Yasuhiro Oda, and Nils Pal Chowdhury for sharing independent observations. This work was supported in part by the US Army Research Office grants W911NF-14-1-0411 and W911NF-17-1-0159, and the National Science Foundation CAREER award MCB-1749489. BEM was also supported by an Indiana Space Grant Consortium fellowship and an IU Dissertation Fellowship.

