## Supplementary material for "Evolved bacterial formate assimilation is likely potentiated by a rudimentary CO_2_ concentrating mechanism": Fig S

for

Breah LaSarre, Department of Plant Pathology, Entomology, and Microbiology, Iowa State University, Ames, Iowa, USA

Alekhya M. Govindaraju, Department of Plant and Microbial Biology, University of California, Berkeley, Berkeley, CA, USA.

**CONTENTS:**

**Monod model for formate-assimilating cultures**

**Single-cell Michaelis-Menten model**

**Fig S1. Formate assimilation pathways**

**Fig S2. ElB RibX repair mutant showed variable growth trends.**

**Fig S3. Comparison of formate versus thiosulfate as electron donors for autotrophic growth where $NaHCO_3$ is used to provide $CO_2$.** Experimental results are in Fig 6A-C.

**Fig S4. Effect of PpsR2 and FbF mutations on *R. palustris* pigmentation**

**Fig S5. Precursor concentration affects CBB activity but population size does not in single cell simulations.**

**Fig S6. Extrapolations of evolved growth rates to time 0**

**Table S1. Complete list of mutations identified in evolved populations and isolates - see separate Excel file**

**Table S2. Strains, plasmids, and primers.**

**Table S3. Default parameters for the Monod Model**

**Table S4. Default parameters for the single-cell model**

**Monod model for formate-assimilating cultures**

Formate-assimilating cultures (Fig 10) were simulated using a Monod model in RStudio (1), that describes:

(1) *R. palustris* growth rate ($h^{-1}$):

$$\mu_{Rp} = \mu_{MAX}\,[CO2/(k_C + CO2)] \cdot [XH/(k_X + XH)] \cdot [For/(k_F + For)]$$

(2) change in $CO_2$ concentration (mM/h):

$$dCO2/dt = \mu_{Rp} \cdot Rp/Y_F - \mu_{Rp} \cdot Rp/Y_C$$

(4) change in formate concentration (mM/h):

$$dF/dt = -\mu_{Rp} \cdot Rp/Y_F$$

(5) change in reductant (e.g., NAD(P)H) concentration (mM/h):

$$dX/dt = \mu_{Rp} \cdot Rp/Y_F - \mu_{Rp} \cdot Rp/Y_X$$

(6) change in *R. palustris* population (cells/ml/h):

$$dRp/dt = \mu_{Rp} \cdot Rp$$

where,

$\mu_{Rp}$ is the specific growth rate ($h^{-1}$).

$\mu_{MAX}$ is the maximum specific growth rate ($h^{-1}$)

CO2 is the $CO_2$ concentration (mM)

XH is the reductant (e.g., NAD(P)H) concentration (mM)

For is the formate concentration (mM)

$k_i$ is the half saturation constant (km) for the indicated substrate (C, $CO_2$; X, XH; F, formate) (mM)

Rp is the *R. palustris* cell density (cells/ml)

Y*i* is the growth yield on the indicated substrate (cells / µmol).

Ff is the formate excretion ratio, which is the same as 1/Yf (µmol / cell)

Initial and default parameter values are shown in Table S2. Simulations assumed that a 0.1 mM pool of reductant (XH) was initially available and that XH could accumulate without consequence. Simulations modified the value of Ff to affect the ratio between the formate oxidation rate and the $CO_2$ fixation rate.

**Single-cell Michaelis-Menten model**

The single cell Michaelis-Menten model described the following:

(1) CBB cycle rate (mM $h^{-1}$):

$$vCBB = vCBB_{MAX} \cdot [CO2/(k_C + CO2)] \cdot [XH/(k_X + XH)]$$

(2) Formate dehydrogenase rate (mM $h^{-1}$):

$$vFDH = vFDH_{MAX} \cdot [For/(k_F + For)] \cdot [1-(XH/(k_X +XH))]$$

(4) Pigment synthesis rate (mM $h^{-1}$):

$$vPig = vPig_{MAX} \cdot [Pre/( k_P + Pre)] \cdot [XH/(k_X + XH)]$$

(5) Carbonic anhydrase rate on $CO_2$ (mM $h^{-1}$):

$$vCAc = Rf \cdot vCA_{MAX} \cdot [pH-(Bic/CO2)] \cdot [CO2/(k_{CA} + CO2)] + Rf \cdot vS_{MAX} \cdot [pH-(Bic/CO2)]$$

(6) Carbonic anhydrase rate on $HCO_3^-$ (mM $h^{-1}$):

$$vCAb = vCA_{MAX} \cdot [pH-(Bic/CO2)] \cdot [BIC/(k_{CA} +BIC)] + vS_{MAX} \cdot [pH-(Bic/CO2)]$$

(7) Spontaneous conversion of $CO_2$ to $HCO_3^-$ (mM $h^{-1}$):

$$vSc = Rf \cdot vS_{MAX} \cdot [8-(Bic/CO2)]$$

(8) Spontaneous conversion of $HCO_3^-$ to $CO_2$ (mM $h^{-1}$):

$$vSb = vS_{MAX} \cdot [8-(Bic/CO2)]$$

(9) change in $CO_2$ concentration (mM $h^{-1}$):

$$dCO2/dt = - vCBB + vFDH + vPig – [(CO2 - Co) \cdot Perm \cdot SA/V] - vCAc + vCAb$$

(10) change in formate concentration; include '+vFDH' to keep formate levels constant by replenishing any formate removed (mM $h^{-1}$):

$$dF/dt = \text{-vFDH} + \text{vFDH}$$

(11) change in reductant concentration (mM $h^{-1}$):

$$dX/dt = \text{-vCBB} + \text{vFDH} – \text{vPig}*10/18$$

(12) change in precursor concentration (mM $h^{-1}$):

$$d\text{Pre}/dt = \text{-vPig}*4/18$$

(13) change in $HCO_3^-$ concentration (mM $h^{-1}$):

$$\text{dBic/dt} = \text{vCAc*CO2} - \text{vCAb*Bic}$$

(14) change in extracellular $CO_2$ concentration (mM $h^{-1}$):

$$\text{dCO2out/dt} = [(\text{CO2} - \text{Co}) \cdot \text{Perm} \cdot \text{SA/V} \cdot \text{Popn/Diln}] – \text{vSc*Co} + \text{vSb*Bo}$$

(15) change in extracellular $HCO_3^-$ concentration (mM $h^{-1}$):

$$\text{dBicout/dt} = \text{vSc*Co} – \text{vSb*Bo}$$

where,

| | |
|---|---|
| vCBB | is the rate of $CO_2$ and XH removal by the Calvin cycle (mM $h^{-1}$) |
| $vCBB_{MAX}$ | is the maximum rate of $CO_2$ and XH removal by the Calvin cycle (mM $h^{-1}$) |
| vFDH | is the rate of $CO_2$ and XH production by FDH (mM $h^{-1}$) |
| $vCBB_{MAX}$ | is the maximum rate of $CO_2$ and XH production by FDH (mM $h^{-1}$) |
| vPig | is the maximum rate of $CO_2$ production from pigment synthesis; when applied to XH removal, it is stoichiometrically scaled by 10/18; when applied to precursor removal it is stoichiometrically scaled by 4/18 (mM $h^{-1}$) |
| $vPig_{MAX}$ | is the maximum rate of $CO_2$ removal from pigment synthesis (mM $h^{-1}$) |
| vCAc | is the conversion rate of intracellular $CO_2$ to $HCO_3^-$ by carbonic anhydrase; Rf is applied to favor an 8:1 ratio of $HCO_3^-$ to $CO_2$ (mM $h^{-1}$) |
| vCAb | is the conversion rate of intracellular $HCO_3^-$ to $CO_2$ by carbonic anhydrase (mM $h^{-1}$) |
| $vCA_{MAX}$ | is the maximum interconversion rate of intracellular $HCO_3^-$ and $CO_2$ by carbonic anhydrase (mM $h^{-1}$) |
| vSc | is the spontaneous conversion of extracellular $CO_2$ to $HCO_3^-$ (mM $h^{-1}$) |

| | |
|---|---|
| vSb | is the spontaneous conversion of extracellular $HCO_3^-$ to $CO_2$ (mM $h^{-1}$) |
| $vS_{MAX}$ | is the maximum spontaneous interconversion rate of extracellular $HCO_3^-$ and $CO_2$ (mM $h^{-1}$) |
| CO2 | is the intracellular $CO_2$ concentration (mM) |
| XH | is the reductant (e.g., NAD(P)H) concentration (mM) |
| For | is the formate concentration (mM) |
| Pre | is the concentration of precursors for pigment biosynthesis (mM) |
| Bic | is the intracellular $HCO_3^-$ concentration (mM) |
| Co | is the extracellular $CO_2$ concentration (mM) |
| Bo | is the extracellular $HCO_3^-$ concentration (mM) |
| $k_i$ | is the half saturation constant (km) for the indicated substrate (C, $CO_2$; X, XH; F, formate; P, precursors; CA, $CO_2$ or $HCO_3^-$) (mM) |
| Rf | is the scaling factor to arrive at the desired ratio of $HCO_3^-$ to $CO_2$ |
| pH | is a factor to limit the ratio of $HCO_3^-$ to $CO_2$; different combinations of Rf and pH were used to arrive at the desired ratio by trial and error |
| Perm | is membrane permeability coefficient for $CO_2$ (cm $h^{-1}$) |
| SA | is the cell surface area ($cm^2$) |
| V | is the cell volume ($cm^3$) |
| Popn | is a scaling factor reflecting the population of cells affecting extracellular concentrations |
| Diln | is a dilution factor applied when $CO_2$ leaves the cell |

180 h was simulated, corresponding to the approximate generation time for the *R. palustris* parent grown with formate and $NaHCO_3$ and initially set $CO_2$ diffusion to zero Initial and default parameter values are shown in Table S3.

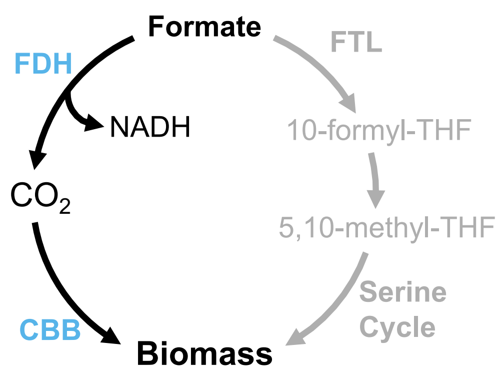


**Fig S1. Formate assimilation pathways.** Grey, *R. palustris* cannot assimilate formate directly via the Serine cycle due to the absence of formate tetrahydrofolate (THF) ligase (FTL). Black, *R. palustris* has genes encoding formate dehydrogenase (FDH) and the CBB cycle, which could allow for indirect formate assimilation via $CO_2$. A challenge with the indirect route is loss of $CO_2$ before it can be captured by Rubisco in the CBB cycle.

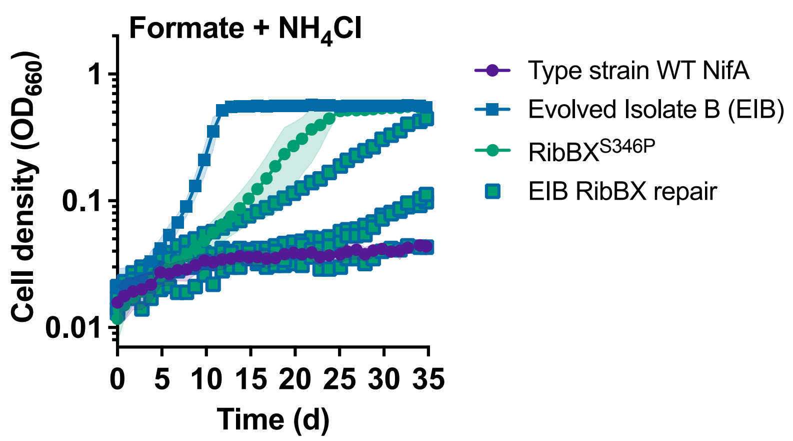


**Fig S2. EIB RibX repair mutant showed variable growth trends.** The figure is the same as Fig 5B but shows all four 'EIB RibBX repair' biological replicates. Shading = SD; n=4. Growth curves for the type strain and evolved isolate B are the same for Fig 5, 8, and 9.

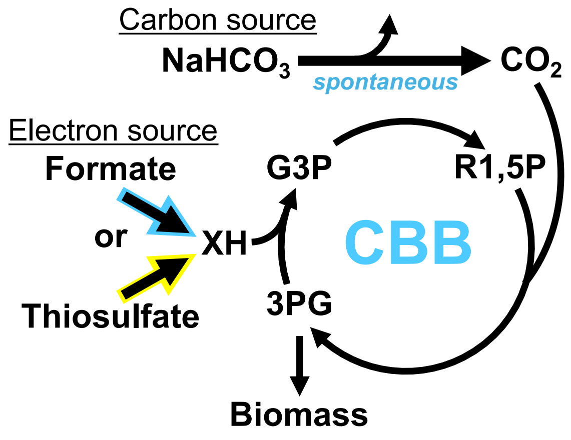


**Fig S3. Comparison of formate versus thiosulfate as electron donors for autotrophic growth where $NaHCO_3$ is used to provide $CO_2$.** Experimental results are in Fig 6A-C.

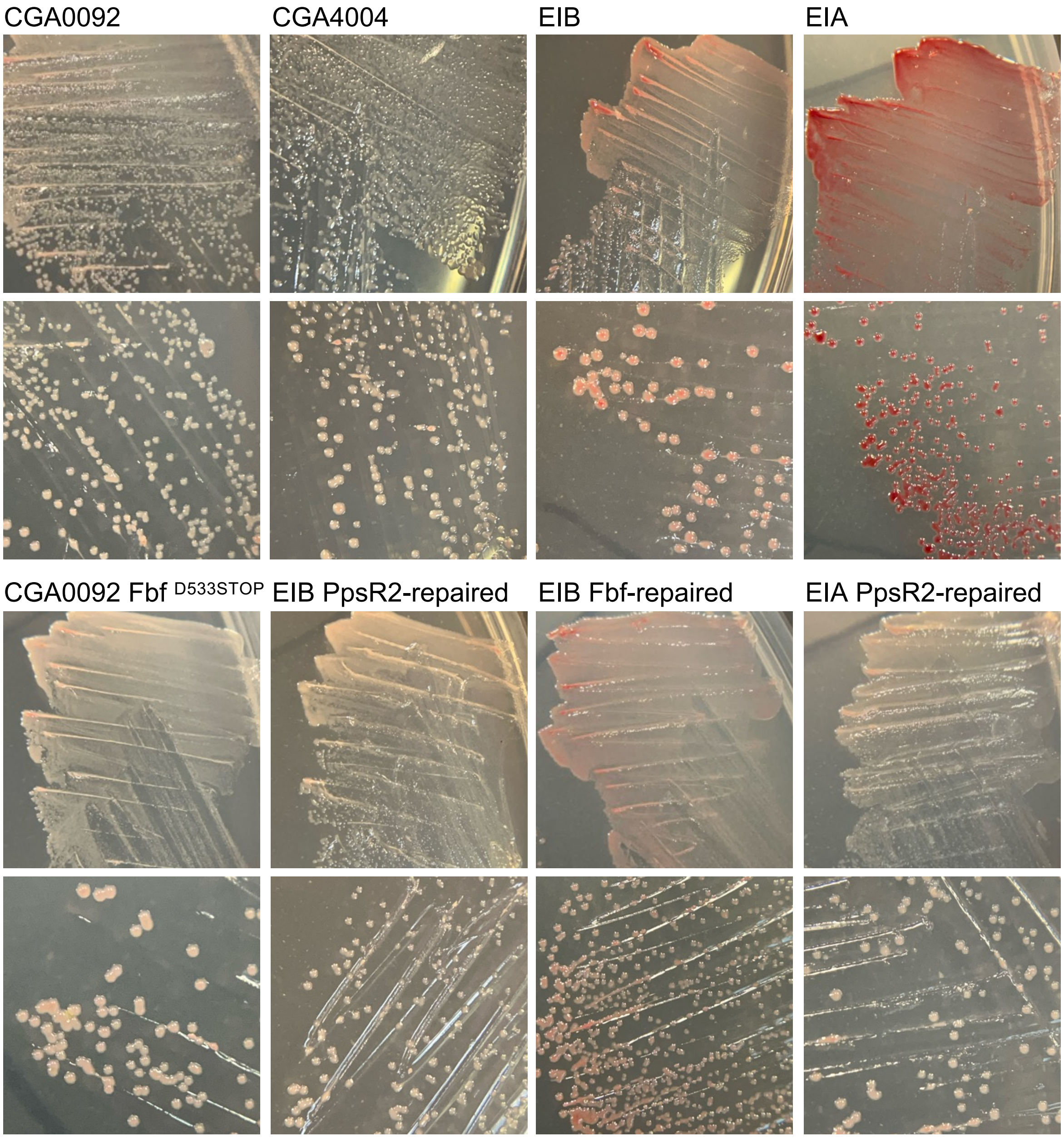


**Fig S4. *ppsR2* mutants are more pigmented than ancestral *R. palustris* strains and than the FbF $^{D533STOP}$ mutant.** All strains were incubated under oxic conditions on PM succinate agar at 30°C in darkness (plates were wrapped in tinfoil) for 7 days. Oxic conditions repress pigment synthesis in the type-strain CGA0092 and the parent strain CGA4004.

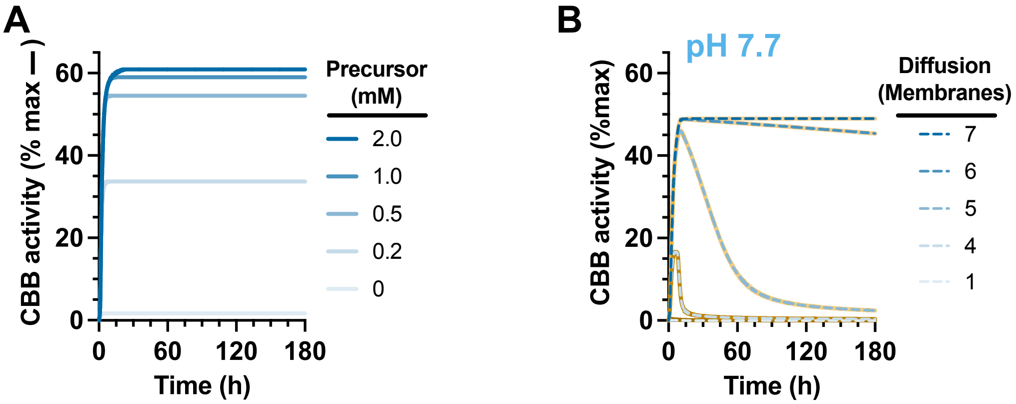


| | | |
|---|---|---|
| Perm coeff (cm $h^{-1}$): | 0 | variable |
| $vPIG_{MAX}$ (mM $h^{-1}$): | 0.64 | 0.64 |
| Precursors (mM): | variable | 1 |

**Fig S5. Precursor concentration affects CBB activity but population size does not in single cell simulations. (A, B)** Single cell Michaelis-Menten model simulations as in Fig 10. **(A)** Effect of pigment precursor concentration **(A)** and membrane permeability based on the number of chromatophore membranes **(B)** on the CBB cycle rate, using the specified parameter values below each graph. **(B)** Orange lines are the same data from Fig 10E, where $CO_2$ production from a population of 1.3 x $10^7$ cells/ml was used. Dashed lines are from simulations where only a single cell was considered, without a population contributing to $CO_2$.

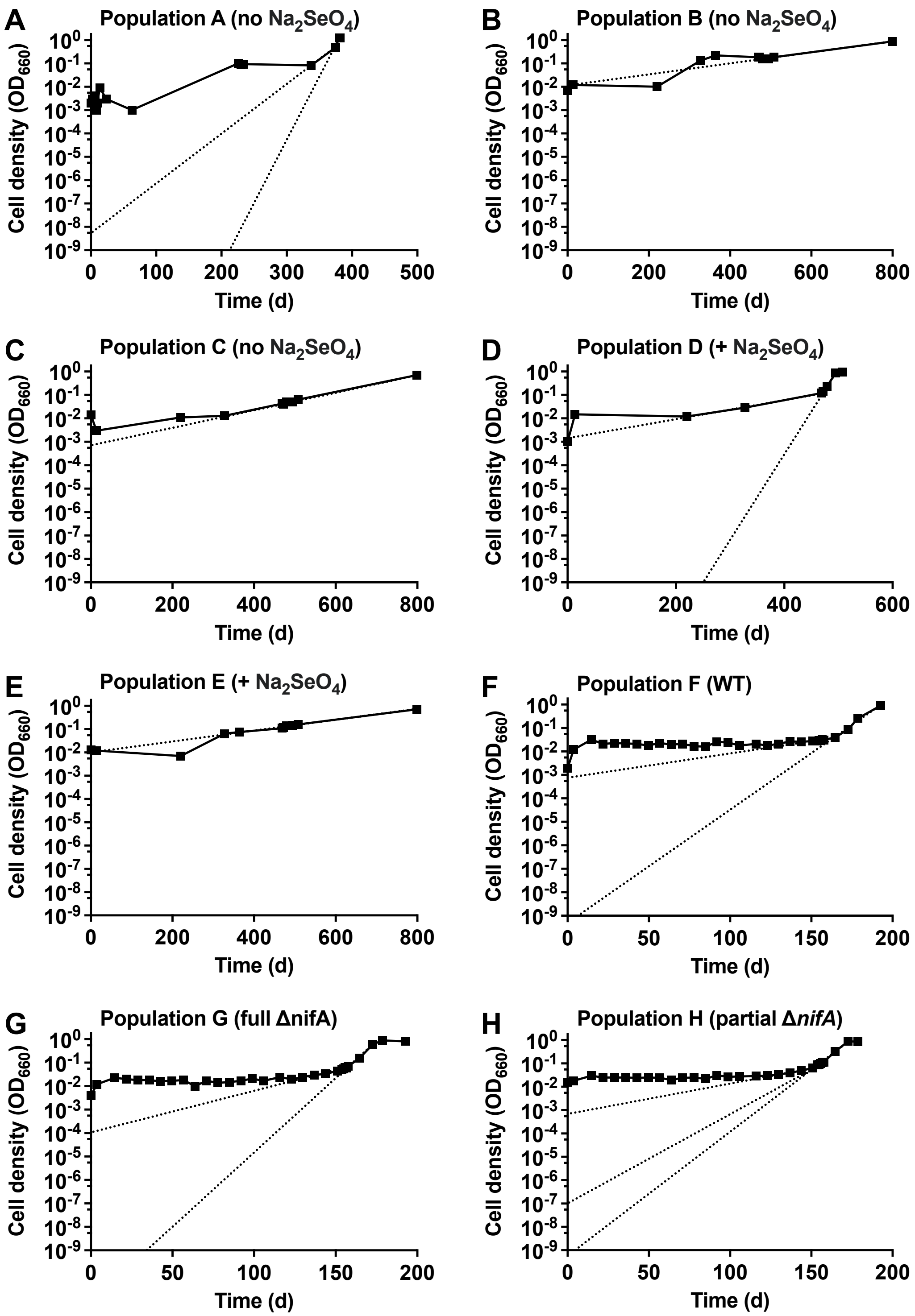


**Fig S6. Extrapolating to estimate initial formate-assimilating subpopulations.** Solid lines and symbols are the same data for T0 cultures from Fig 1 and 4. Dotted lines extrapolate backwards in time from various growth rates observed in each culture. An 1 $OD_{660}$ roughly corresponds to $10^9$ cells per ml, thus we use $10^{-9}$ $OD_{660}$ here to represent 1 cell / ml.

**Table S2. Strains, plasmids, and primers.**

| Strain/Plasmid/ Primer, Designation | Genotype/Plasmid use/Sequence | Source/Primer use |
|---|---|---|
| *Rhodopseudomonas palustris* | | |
| CGA0092, Type strain | Wild type | (2) |
| CGA4004, Parent | CGA0092, Δ*uppE,* Δ*hupS* | (3) |
| CGA4005, Ancestor | CGA0092, *nifA*,* Δ*uppE,* Δ*hupS* | (3) |
| CGA4028 | CGA0092, full Δ*nifA* | This study |
| CGA757-2 | CGA0092, partial Δ*nifA* (1,332 bp deleted) | This study |
| CGA4005 Evolved Isolate A | Table S1 | This study |
| CGA4005 Evolved Isolate B, EIB | Table S1 | This study |
| CGA4005 Evolved Isolate C | Table S1 | This study |
| CGA4005 Evolved Isolate D | Table S1 | This study |
| CGA4005 Evolved Isolate E | Table S1 | This study |
| | CGA0092 $RibBX^{S336P}$ | This study |
| | CGA4004 $RibBX^{S336P}$ | This study |
| | Evolved Isolate B RibBX repair | This study |
| | CGA0092 $PpsR2^{I402N}$ | This study |
| | Evolved Isolate B PpsR2 repair | This study |
| | CGA0092 $Fbf^{D533STOP}$ | This study |
| | Evolved Isolate B Fbf repair | This study |
| | CGA4004 NifAL563R | This study |
| | CGA4005 NifAL563R | This study |
| | Evolved Isolate B Δ*cbbP*::KmR | This study |
| | Evolved Isolate B Δ*fdsGBA*::KmR | This study |
| *Escherichia coli* | | |
| NEB10β | DH10B derivative; *Δ(ara-leu) 7697 araD139 fhuA ΔlacX74 galK16 galE15 e14- ϕ80dlacZΔM15 recA1 relA1 endA1 nupG rpsL* ($Str^R$) *rph spoT1 Δ(mrr-hsdRMS-mcrBC)* | New England Biolabs |
| **Plasmids** | | |
| pJQ200SK | *R. palustris* suicide vector | (4) |
| $pJQNifA^{L563R}$ | Suicide vector to introduce $NifA^{L563R}$ | This study |
| $pJQRibBX^{S336P}$ | Suicide vector to introduce $RibBX^{S336P}$ | This study |
| $pJQPpsR2^{I402N}$ | Suicide vector to introduce $PpsR2^{I402N}$ | This study |
| $pJQFbf^{D533STOP}$ | Suicide vector to introduce $Fbf^{D533STOP}$ | This study |
| pJQWTRibBX | Suicide vector to introduce the WT *ribB* allele | This study |
| pJQWTPpsR2 | Suicide vector to introduce the WT *ppSR2* allele | This study |
| pJQWTFbf | Suicide vector to introduce the WT *fbf* allele | This study |
| pJQΔ*nifA* | Suicide vector to introduce the full Δ*nifA* | This study |
| pJQΔ*nifA-1332* | Suicide vector to introduce the partial Δ*nifA* | (5) |

| | | |
|---|---|---|
| pJQΔ*cbbP*::Km$^R$ | Suicide vector to introduce Δ*cbbP*::Km$^R$ | (6) |
| pJQΔ*fdsGBA*::Km$^R$ | Suicide vector to introduce Δ*fdsGBA*::Km$^R$ | This study |
| pBBPgdh | Empty vector | (7) |
| pBBP*cbbP* | *cbbP* complementation vector | (6) |
| pPS858 | Source of the kanamycin resistance cassette | (7) |
| Primers (Forward, F; Reverse, R) | | |
| BEH41 | caaaagctggagctccaccgCACGCAGGTGTTGTCGAC | F; pJQNifA$^{L563R}$ assembly |
| BEH42 | atcgaattcctgcagcccggATGTAGTGCCGATGATCCTG | R; pJQNifA$^{L563R}$ assembly |
| BEH70 | gcgctggatctgctgaagg | F; NifA$^{L563R}$ screening |
| BEH71 | gcttgtcgaactgaccttcgca | R; NifA$^{L563R}$ screening |
| BEH59 | gcgaaccccagagtccc | F; Km$^R$ cassette amplification from pPS858 |
| BEH60 | tccgctagcttcacgct | R; Km$^R$ cassette amplification from pPS858 |
| BEH93 | caaaagctggagctccaccgaacatgcagaatcgccgc | F; Δ*fds* left fragment |
| BEH94 | agcgggactctggggttcgcttcgtattgcatcggcgc | R; Δ*fds* left fragment |
| BEH95 | ggcagcgtgaagctagcggagaatacaaggtcacggcc | F; Δ*fds* right fragment |
| BEH96 | atcgaattcctgcagcccggacatcctcacgcacatgc | R; Δ*fds* right fragment |
| BEH97 | gatggttcgtgcttccc | F; Δ*fdsGBA*::Km$^R$ screening |
| BEH98 | ccaacgagcttgtcgag | R; Δ*fdsGBA*::Km$^R$ screening |
| BEH99 | ggatcgaacttcaggaagc | F; Δ*cbbP*::Km$^R$ screening |
| BEH100 | atgtcgttcgtgatggag | R; Δ*cbbP*::Km$^R$ screening |
| BEH119 | caaaagctggagctccaccgGAGCTCagaactcgatctccagcg | F; Fbf$^{D533STOP}$ assembly |
| BEH120 | atcgaattcctgcagcccggCTCGAGtcaagctggtggcattcc | R; Fbf$^{D533STOP}$ assembly |
| BEH121 | agcaacagagacaggaccagc | F; Fbf screening |
| BEH122 | ttcttcggcgggctgatttg | R; Fbf screening |
| BEH123 | caaaagctggagctccaccggtcctgtttcagagatcc | F; RibBX$^{S336P}$ assembly |
| BEH124 | atcgaattcctgcagcccggattgaactgcttgcggtc | R; RibBX$^{S336P}$ assembly |
| BEH125 | gttccacaagccgaacatcgt | F; RibBX screening |
| BEH126 | ctcgtcttcggtgagctgatctt | R; RibBX screening |
| BEH135 | gatctgtccagcttgctgac | F; PpsR2 screening |
| BEH136 | cttcacgttcgcaattcgaaag | R; PpsR2 screening |
| BEH137 | gggaacaaaagctggagctccaccggcatctcggctttccgcc | F;PpsR2$^{I402N}$ assembly |
| BEH138 | ttgatatcgaattcctgcagcccgggcagataatggtgacaggcaac | R;PpsR2$^{I402N}$ assembly |
| BL515 | GACTCTCGAGtgcatgcgctgatcaacaatgcg | F; Δ*nifA* upstream, XhoI |
| BL516 | GACTGGTACCagccatagctggtctccatc | R; Δ*nifA* upstream, KpnI |
| BL517 | GACTGGTACCttctgacccttctcgcttgaaatc | F; Δ*nifA* downstream, KpnI |
| BL518 | GACTCTCGAGgatgattcttcaccttctcccaga | R; Δ*nifA* downstream, XhoI |

**Table S3. Default parameters used to simulate formate-assimilating cultures.**

| Parameter | Value | Description (Units); Source |
|---|---|---|
| $\mu_{MAX}$ | 0.019 | *R. palustris* max growth rate observed by EIB on formate alone ($h^{-1}$); |
| CO2 | $1 \times 10^{-6}$ | Initial $CO_2$ (mM); $2 \times 10^{-2}$ mM was used for some simulations, estimated based on carry over from starter cultures (8) |
| XH | 0.1 | Initial XH (mM) |
| For | 20 | Initial formate (mM) |
| Rp | $1.3 \times 10^{7}$ | Initial *R. palustris* cell density (cells / ml) |
| $Y_C$ | $2.8 \times 10^{7}$ | growth yield on formate with $NaHCO_3$ available (electron-limited) (cells / μmol); Fig 6A, assumes all formate was assimilated via $CO_2$ and $1 \times 10^{9}$ cells/ml/$OD_{660}$ |
| $Y_X$ | $1.12 \times 10^{8}$ | growth yield per NAD(P)H oxidized (cells / μmol); assumes a net of 14,270 μmol XH oxidized per cell based on (7) |
| $Y_F$ | $2.8 \times 10^{7}$ | growth yield on formate with $NaHCO_3$ available (electron-limited) (cells / μmol); Fig 6A, assumes all formate was assimilated and $1 \times 10^{9}$ cells/ml/$OD_{660}$ |
| $k_C$ | 0.067 | Rubisco half-saturation constant (Km) for $CO_2$ (mM); (9) |
| $k_X$ | 0.0001 | Set arbitrarily low to assume saturating availability |
| $k_F$ | 0.26 | FDH half-saturation constant (Km) for formate (mM); (10) |

Some values were found via BRENDA (11), BioNumbers (12), and Biocyc (13).

**Table S4. Default parameters used to in the single cell model.**

| Parameter | Value | Description (Units); Source |
|---|---|---|
| $vCBB_{MAX}$ | 938 | Maximum CBB cycle rate, 1 cell ÷ growth yield on $CO_2$ (cells/ml /mM $CO_2$) x growth rate of the parent strain on formate with $NaHCO_3$ ($h^{-1}$) = mM $h^{-1}$; Fig 6A |
| $vFDH_{MAX}$ | 1618 | Maximum FDH rate, assumed to be the same as $vCBB_{MAX}$ (mM $h^{-1}$) |
| $vPig_{MAX}$ | 0.640 (EIB)<br>0.466 (EIB)<br>0.093 (WT) | Maximum $CO_2$ production rate from pigment synthesis, $CO_2$ from BChl (mM; 1 cell x 5 x $10^{20}$ mol BChl/cell (14) x 10-times more BChl in anaerobic phototrophs than aerobic x 18 CO2 per BChl) + $CO_2$ from carotenoids (mM; BChl value x 0.64 caroteinoids/BChl (15) x 4/18 $CO_2$) x growth rate of the strain of interest on formate with $NaHCO_3$ ($h^{-1}$) = mM $h^{-1}$; 0.466 mM $h^{-1}$ applies to EIB without $NaHCO_3$ |
| $vCA_{MAX}$ | 6.12 x $10^9$ | Maximum carbonic anhydrase rate; *Neisseria gonorrhoeae* ($h^{-1}$); (16) |
| $vS_{MAX}$ | 130 | ($h^{-1}$); (17) |
| $CO_2$ | 1 x $10^{-7}$ | Initial intracellular $CO_2$ concentration (mM) |
| XH | 2 | Initial intracellular XH concentration (mM) |
| For | 10 | Initial intracellular formate concentration (mM) |
| Pre | 1 | Initial intracellular pigment precursor concentration (mM) |
| Bic | 8 x $10^{-7}$ | Initial intracellular $HCO_3^-$ concentration (mM) |
| Co | 1 x $10^{-7}$ | Initial extracellular $CO_2$ concentration (mM) |
| Bo | 8 x $10^{-7}$ | Initial extracellular $HCO_3^-$ concentration (mM) |
| $k_C$ | 0.067 | Rubisco half-saturation constant (Km) for $CO_2$ (mM); (9) |
| $k_X$ | 0.01 | Assumed (mM) |
| $k_F$ | 0.26 | FDH half-saturation constant (Km) for formate (mM); (10) |
| $k_P$ | 0.01 | Assumed (mM) |
| $k_{CA}$ | 20 | from *Neisseria gonorrhoeae* (18) |
| Rf | 100 | scaling factor to arrive at an ~8:1 ratio of $HCO_3^-$ to $CO_2$ |
| | 1200 | scaling factor to arrive at a ~40:1 ratio of $HCO_3^-$ to $CO_2$ |
| pH | 8 | used for a ~8:1 ratio of $HCO_3^-$ to $CO_2$ |
| | 1000 | used for a ~40:1 ratio of $HCO_3^-$ to $CO_2$ |
| Perm | | (cm $h^{-1}$) |
|  | 360 | MDCK cells and cholesterol-rich vesicles; (19) |
|  | 36 | carboxysome; (20) |
| | | For chromatophore membranes, we assumed 18-carbon fatty acids (21), that each membrane decreases the permeability coefficient by a factor of 0.037 (1.5-fold decrease for every 2 carbons (22)), and that 1 membrane layer has a permeability of 360 cm $h^{-1}$ such that: |
| | 6.7 x $10^{-4}$ | 4 membranes |
| | 2.5 x $10^{-5}$ | 5 membranes |
| | 9.2 x $10^{-7}$ | 6 membranes |
| | 3.4 x $10^{-8}$ | 7 membranes |
| SA | 5.1 x $10^{-8}$ | Surface area for a 3.25 μm long *R. palustris* cell ($cm^2$); (23) |
| V | 6.05 x $10^{-13}$ | Volume for a 3.25 μm long *R. palustris* cell ($cm^3$); (23) |
| Popn | 13 x $10^6$ | Typical initial population of *R. palustris* cells (cells/ml) |
| Diln | 1.65 x $10^{12}$ | Dilution factor going from intracellular volume to 1 ml culture |

Some values were found via BRENDA (11), BioNumbers (12), and Biocyc (13).
